# Optimizing effective labeling efficiency in MINFLUX 3D DNA-PAINT microscopy by maximizing marker detection probability

**DOI:** 10.1101/2025.05.16.654501

**Authors:** Christian Soeller, Alexandre F. E. Bokhobza, Javier Casares-Arias, Alexander H. Clowsley

## Abstract

MINFLUX is a powerful single-molecule approach capable of achieving high spatially isotropic resolution in three dimensions. Current implementations collect localizations strictly serially, but criteria for when to terminate acquisition are often unclear. We therefore systematically investigate the time course of effective labeling efficiency (ELE) and achievable saturation values in MINFLUX 3D DNA-PAINT microscopy of Nup96 proteins in a U-2 OS-Nup96-mEGFP modified cell line using a commercial MINFLUX microscope. ELE was measured with a maximum-likelihood template fitting assisted quantitative procedure. We collected data measured over various scan sizes and achieved ELE values of ∼60% after passing a time interval dependent on the region size, typically requiring long-duration acquisitions over several hours. Our data and a simple model suggest that maximizing marker detection is key to achieving the limits set by chemical labeling efficiency. A factor limiting the marker detection probability when using conventional DNA-PAINT markers is docking strand site-loss, observed over the duration required to build up the image data of MINFLUX acquisitions, which also limits the achievable number of labeling site visits to values around 1-3. Using repeat DNA-PAINT, i.e. employing oligonucleotide sequences with repeated docking sites, we observed greatly reduced site-loss and could increase the number of individual visits to site locations by more than threefold over the same period. Additionally, this enabled increasing stringency criteria for labeling (i.e. higher threshold values) and maximizing marker detection probabilities so that ELE reaches the limits set by chemical labeling efficiency.

## Background

MINFLUX ‘nanoscopy’ is a second-generation super-resolution approach capable of obtaining nanometer precision in three dimensions.^1^ The technique builds upon the foundation of the original super-resolution approaches, STED^2,3^ and STORM/PALM,^4,5^ by combining elements of both. MINFLUX systems rely on rapidly scanning an excitation beam with a known intensity minimum, such as a donut in 2D and a bottle beam pattern for 3D localization, Supp. Figure 1a & 1b, across a sample and attempts to capture single-molecule fluorescent signals within the central zero position. One can imagine that, as the beam closes in on its target fewer photons are collected, and the known position of the excitation intensity zero is used to estimate the position of the fluorophore, Supp. Figure 1c.

Generally, in single-molecule localization microscopy (SMLM), the generation of individual events has typically been achieved by using photo-switching fluorophores such as Alexa Fluor 647, which require special chemical switching buffers.^6^ These dyes are generally permanently attached to the marker being imaged and, as such, eventually photo-bleach into an unexcitable state. To overcome this limitation, approaches using complementary strands of DNA with photostable dyes are becoming more common, termed DNA-PAINT,^7^ and are also being applied to MINFLUX.^8,9^ DNA-PAINT uses dye-modified oligonucleotide sequences, free in-solution, called ‘imagers’, which are complementary to ‘docking’ strand sites affixed to the target marker.^7,10^ These imagers are free to stochastically, and importantly, reversibly hybridize to an available docking site with a designed mean binding time that is typically a fraction of a second. It is this transient immobilization of the imager that culminates in a detectable single-molecule event. DNA-PAINT, with appropriate excitation parameters and dye choices, circumvents the effects of photobleaching due to the constant replenishment of imagers present within the buffer, however, an alternative loss mechanism termed site-loss has been shown to occur from phototoxicity.^11,12^ In this site-loss scenario, docking strands become chemically modified and accordingly are unable to bind imager stands with the original designed affinity and attachment time. Nevertheless, the versatile approach has enabled the construction of a number of DNA-nanotools for biological investigation at the super-resolution level^13,14^ and innovations have been demonstrated to combat non-specific interactions in biological samples from both imager^15^ and marker labeling.^16^

Some previous publications have suggested that MINFLUX microscopy may sometimes undersample known structures^17,18^ while some disagreement in the literature exists^19^. This discussion highlights that MINFLUX effective labeling efficiency has, to our knowledge, not been previously systematically quantified. This contrasts with conventional wide-field super-resolution microscopy where proteins in the nuclear pore complex have been introduced as a suitable reference sample. Quantitative procedures to determine the effective labeling efficiency were established in that study, suggesting values between 40-80% can be consistently achieved with suitable high-quality marker systems^20^.

We sought to fill this gap by translating the use of NPC labeling and imaging of Nup96-mEGFP distribution to 3D MINFLUX microscopy to test for potential differences in achievable effective labeling efficiency as compared to widefield super-resolution microscopy. Our 3D NPC-based labeling efficiency assays show that MINFLUX DNA-PAINT can achieve similar labeling efficiencies as observed in widefield super-resolution microscopy, but only if care is taken to image for extended durations. As compared to widefield DNA-PAINT, site-loss seems more pronounced in MINFLUX DNA-PAINT but is still compatible with achieving effective labeling efficiencies >50%. We demonstrate that site-loss can be effectively mitigated through the use of repeat docking domains, enabling DNA-PAINT experiments to run unaided for hours^12^ thus fully attaining the labeling efficiency limits set by chemical constraints that generally only depend on the marker system in use and the accessibility of the target sites. Anecdotal use of dSTORM-based dye switching using labile switching buffers was, at least in our hands, more problematic in achieving high labeling efficiencies and may have contributed to the impression of undersampling noticed on occasion^17,18^.

## Results

### Effective labeling efficiency (ELE)

Achieving a high labeling density is critical in super-resolution imaging. In fluorescence microscopy, specifically SMLM, achieving a high effective labeling efficiency involves two major contributions. There is an initial probability of successfully labeling each possible site based on the chemical labeling efficiency, p_chem_, which is influenced by several factors including the density of probes and their accessibility to the labeling target etc. Once the sample is appropriately labeled to be imaged on a super-resolution setup there is a further contribution from the detection probability, p_detect_. For a MINFLUX microscope this culminates in the probability of detecting the marker; a stochastic single-molecule event, coinciding with the scan location, overcoming the acquisition detection criteria (for MINFLUX microscopy these criteria are implicit in the acquisition “sequence” definition, see Methods).

In SMLM and specifically in MINFLUX microscopy, many images and/or events are sequentially detected over an extended time, accordingly p_detect_ generally increases with imaging time. Eventually, a limiting maximal value is reached that in DNA-PAINT may be influenced by site-loss/marker damage by reactive oxygen species photo-damage. Together, the labeling and detection probabilities determine the effective labeling efficiency *P*_*ELE*_ = p_chem_p_detect_. Here we use a prototypical labeling system, using by now well-established single-domain antibodies (sdAB) against GFP, and modified with docking strands for DNA-PAINT, to label Nup96-eGFP in a U-2 OS cell line and investigate how to maximize the detection probability in MINFLUX imaging, while having an essentially unaltered chemical labeling efficiency, p_chem_, across experiments. Using the NPC-based labeling system we measure the effective labeling efficiency with MINFLUX DNA-PAINT in 3D while varying imaging parameters and identify conditions to maximize p_detect_ with the goal that *P*_*ELE*_ should approach p_chem_ in the best-case scenario.

### Imaging of the nuclear pore complex (NPC) as a convenient biological standard

We opted to use the CRISPR-engineered U-2 OS cell line endogenously expressing GFP on Nup96 proteins as they provide a well-characterized biological structure with, effectively, a well-known protein arrangement.^20^ With a diffraction limited confocal scan of the MINFLUX microscope, the Nup96 protein locations are not resolved, Figure 1a. Instead, one sees a collection of essentially featureless puncta, where each punctum of GFP signal encompasses the position of a whole NPC, enabling a general initial assessment of the nature of the cell (inset panel Figure 1a). Only once super-resolved using DNA-PAINT (see “blinks” recorded in the confocal mode of the MINFLUX microscope, Fig. 1bi) do Nup96 locations become more apparent, as can be seen in two views of a single NPC structure recorded as part of an extended duration MINFLUX acquisition (Fig. 1bii, iii).

**Figure 1.**
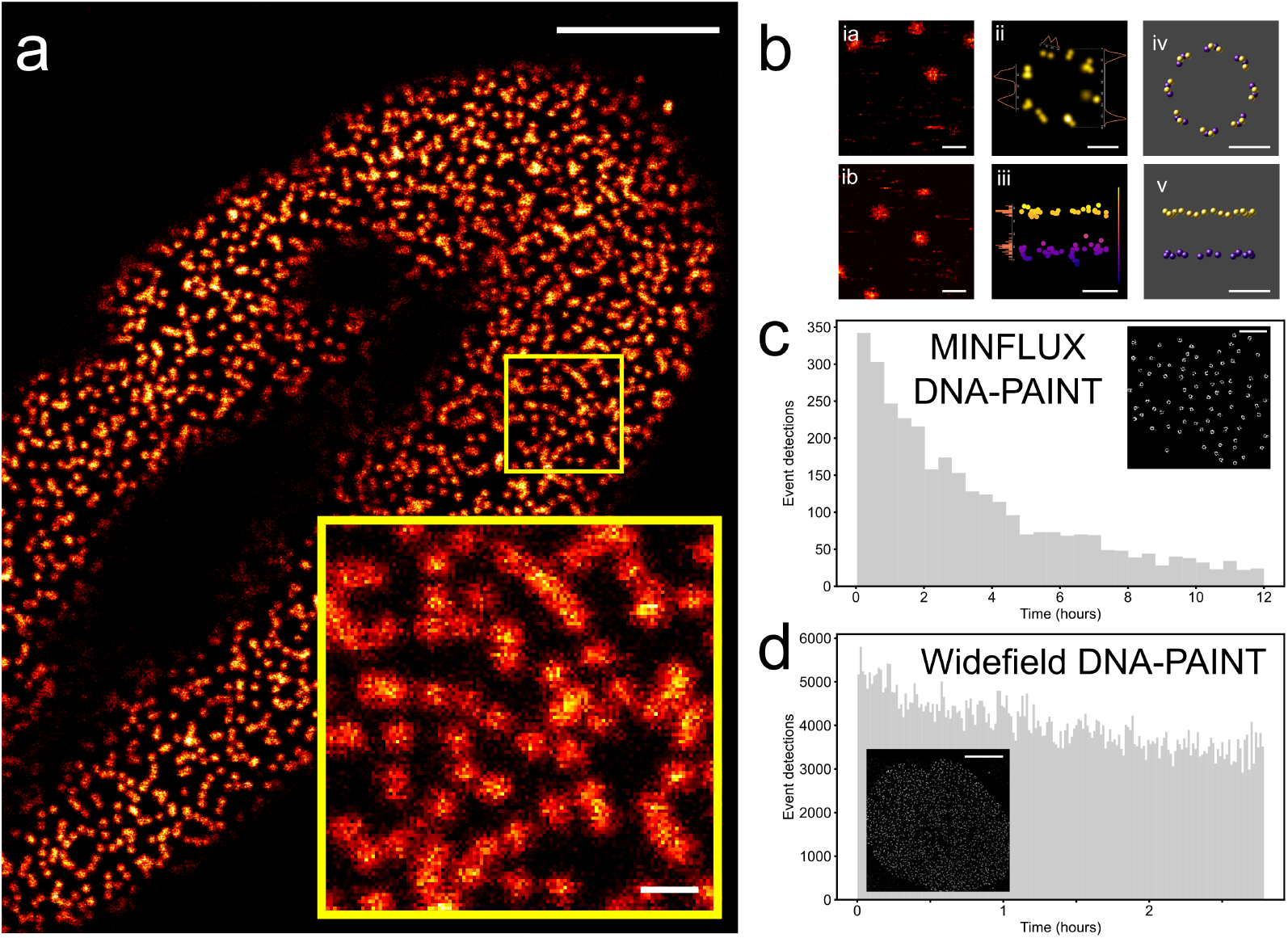
SMLM super-resolution imaging of NPC proteins. a. A confocal overview image of a U-2 OS-Nup96-mEGFP modified cell. The zoomed-in image of the ∼3 x 3 µm^2^ boxed region shows the mEGFP diffraction-limited signal displaying the relative position of nuclear pore complexes (NPCs). **b**. i a,b: Frames from fast confocal scans showing the transient attachment of imagers to markers on Nup96-mEGFP; ii, iii: x-y and x-z views of MINFLUX localizations associated with a single NPC structure; iv, v: schematic of the arrangement of Nup96-mEGFP targets. **c**. The sequential scanning applied in MINFLUX emphasizes the effects of docking strand site damage with a reduction in the number of events detected over the long time needed to build up NPC structures. The inset shows an example rendered image. **d**. In widefield DNA-PAINT imaging, acquisition times are briefer to sample all NPCs in the field of view. Accordingly, event rates remain largely undiminished, inset shows an example rendered image. Scale bars: **a**: 5 µm, inset: 500 nm, b ia&ib: 500 nm & ii-iv: 50 nm, **c**: 1 µm, **d**: 5 µm.

The MINFLUX localizations reflect the structure of the Nup96 site distribution, where, in the plane of the nuclear envelope, individual Nup96 proteins are spaced ∼12 nm apart between pairs and each separated ∼42 nm from their closest neighboring pair making a symmetrical ring of eight, as shown in the schematic Fig. 1b iv. This pattern is replicated on both sides of the nuclear envelope to give essentially two “rings” on both cytoplasmic and nucleoplasmic sides separated by ∼50 nm (Fig. 1b v). In total, 32 Nup96 proteins can be labeled per NPC.

In MINFLUX 3D DNA-PAINT we observe a fairly pronounced decay of localization rates over the time course required to build up the structure (Fig. 1c). The time required increases as compared to widefield acquisition due to the entirely serial nature of the acquisition and the decay of the localization rate appears to result from site-loss as we show further below. By contrast (Fig. 1d), an overview of a widefield DNA-PAINT image of Nup96 signals is shown together with the time course of localizations that exhibits a much more gradual decay over time scales that fully build up the structure (typically <1h, Supp. Fig. 2). To quantify these qualitative observations, we established a refined version of the approaches in Thevathasan et al.^20^ to more precisely measure effective labeling efficiency, here specifically tailored to the 3D MINFLUX microscopy modality.

**Figure 2.**
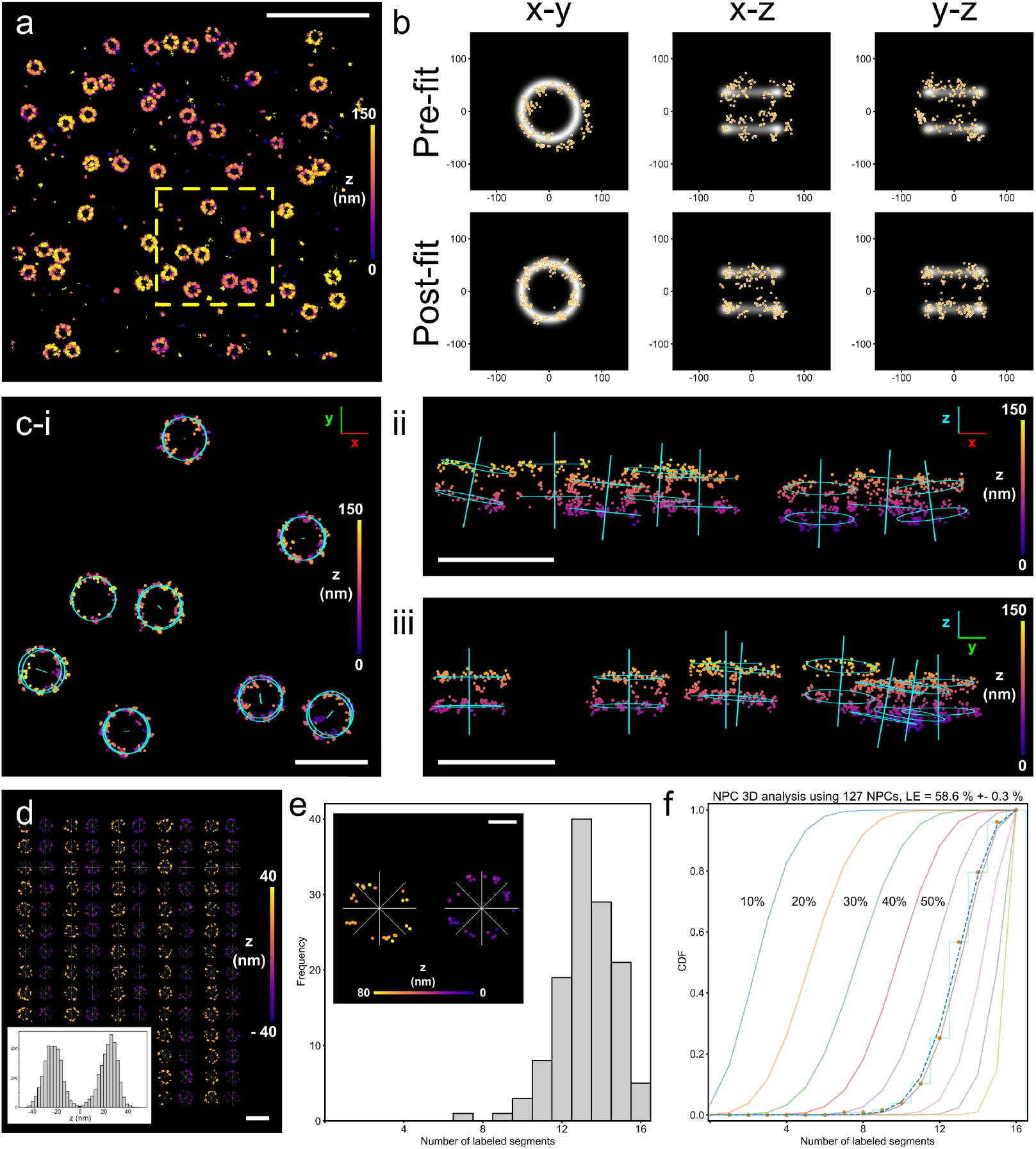
3D NPC labeling effciency methodology. **a)** A typical region of U-2 OS-NUP96-mEGFP NPCs localized using a MINFLUX microscope. **b)** Individual NPC localizations, shown in x-y, x-z and y-z, displayed over a double ringed template structure for both pre-fit and post-fit localizations whereby transformations were applied to achieve the best fit with the template. **c-i)** Fitted NPCs, taken from the boxed region in **a**, show the additional geometry from the points fitted to the rings in x-y. Axial visualization, **c-ii**: x-z, and **c-iii**: y-z demonstrate the subtle curvatures of the nuclear envelope, indicated by the axis along the pore which was added to the template overlay for clarity. **d)** A gallery representation of the nucleoplasmic and cytoplasmic rings, for individual NPCs, displayed next to one another enables a quick assessment of the quality of the data. A histogram of these NPCs exhibits an approximate 50 nm peak to peak separation in z. **e)** A histogram showing the number of NPC segments labeled for the dataset. The localizations are separated into 8 segments for both top and bottom NPC rings, see inset. **f)** Cumulative distribution function plot of labeled segments (data, red dots) and estimated effective labeling effciencies based on the model fit (dashed line). Solid lines show model predictions for 10-90% ELE. Color maps indicate localized position in z. Scale bars: a: 1 µm, c-i, c-ii & c-iii: 200 nm, d & e: 50 nm.

### MINFLUX DNA-PAINT 3D imaging of NPCs and quantification of ELE

#### Titrating Imager concentrations for DNA-PAINT

The rate of single-molecule events in DNA-PAINT is dependent on several factors, including: the number of available docking sites on a marker, the density of the markers, the imager concentration, and the buffer composition. Prior to MINFLUX experiments, we therefore took short *xyt* confocal scans with set parameters (see also Fig. 1bi and Methods), to ascertain appropriate imager blink rates. These scans enabled us to semi-quantitatively assess the level of imager to docking-site bindings and reduce the likelihood of overlapping events by titrating the imager concentration, see also Supp. Media 1. This is especially important in MINFLUX microscopy as such overlaps can lead to the failure of MINFLUX localizations and/or localization errors^21^. Conversely, the imager concentrations should be high enough to achieve a high rate of successful localizations to keep acquisition times in a reasonable range.

Prior to NPC labeling analysis, localizations from the same trace were coalesced to a single detection at the centroid of the ‘trace’ based on the inherent ‘traceID’ property, with lower localization error resulting from coalescing, as described previously^9^. All events in the same trace will originate from a single imager binding to a single docking strand and should therefore be treated as a single “compound localization”.

#### Maximum-likelihood fitting of NPC structures

To estimate ELE, we used an approach similar to the method introduced by Thevathasan et al.^20^, but extended to 3D for MINFLUX microscopy. MINFLUX 3D data of Nup96-mEGFP labeled with anti-GFP DNA-PAINT sdABs (Fig. 2a) were rendered to a 2D image in PYMEvisualize and locations of NPCs were detected by a semi-manual approach in Fiji^22^ and stored as regions-of-interest (ROIs) in ImageJ ROI format. ROIs were read into the PYMEvisualize^23^ SMLM data visualizer and used to label coalesced 3D MINFLUX localizations belonging to identified NPCs with a unique ID. We then used maximum-likelihood fitting to a “double ring” template to align NPC events with the template using an algorithm as described^24^ which we implemented into PYMEvisualize^23^. Following maximum-likelihood fitting, NPC events were generally well-aligned with the template (Fig. 2b). The transformed coordinates are used for further determination of labeling of NPC segments in the two rings. In addition, the optimal fitting parameters are also used to display a suitably rotated template schematic overlaid with the original MINFLUX localizations in PYMEvisualize, see Fig. 2 c-i,ii,iii. In addition to the two rings, the overlays contain the central symmetry axis of each NPC structure and reveal the variation in NPC orientation across the field of view, presumably reflecting variations in the plane of the nuclear envelope.

#### Histograms and fitting with a model prediction

Fig. 2d shows a “gallery” of all fitted NPC structures from one MINFLUX 3D dataset, separated into cytoplasmic (blue/violet) and nucleoplasmic (orange/yellow) components, side by side, displaying the axial separation of the contribution from the two sides at the expected distance of ∼50 nm (Fig. 2d, inset). After alignment to “segment boundaries” all fitted NPC localization datasets are also rotationally aligned (Fig. 2e, inset) and for each segment, separately for cytoplasmic and nucleoplasmic sides, the segments were checked if labeled using a selectable threshold (typically a minimum of 1 or 3 events per segment, see also threshold determination in Supp. Fig. 3). This was used to construct histograms (Fig. 2e) showing for the NPC dataset the number of segments labeled (from 0 to 16). The histograms were converted to cumulative form and, in this form, plotted against the predicted model behavior (showing model curves for 10 to 90 % labeling as solid curves), with the model constructed based on the assumption of independent labeling at all 32 Nup96 sites (see supplementary methods). A value of best fit to the model was then determined. The procedure thus culminates in providing this value of best fit, in the sample shown in Fig. 2f obtaining a value of *P*_*ELE*_ = 58.6 ± 0.3 % for a large ROI dataset containing 127 NPC structures. We proceeded to use this approach to determine the time course and limiting value of *P*_*ELE*_ for different MINFLUX region sizes using conventional DNA-PAINT docking strands.

**Figure 3.**
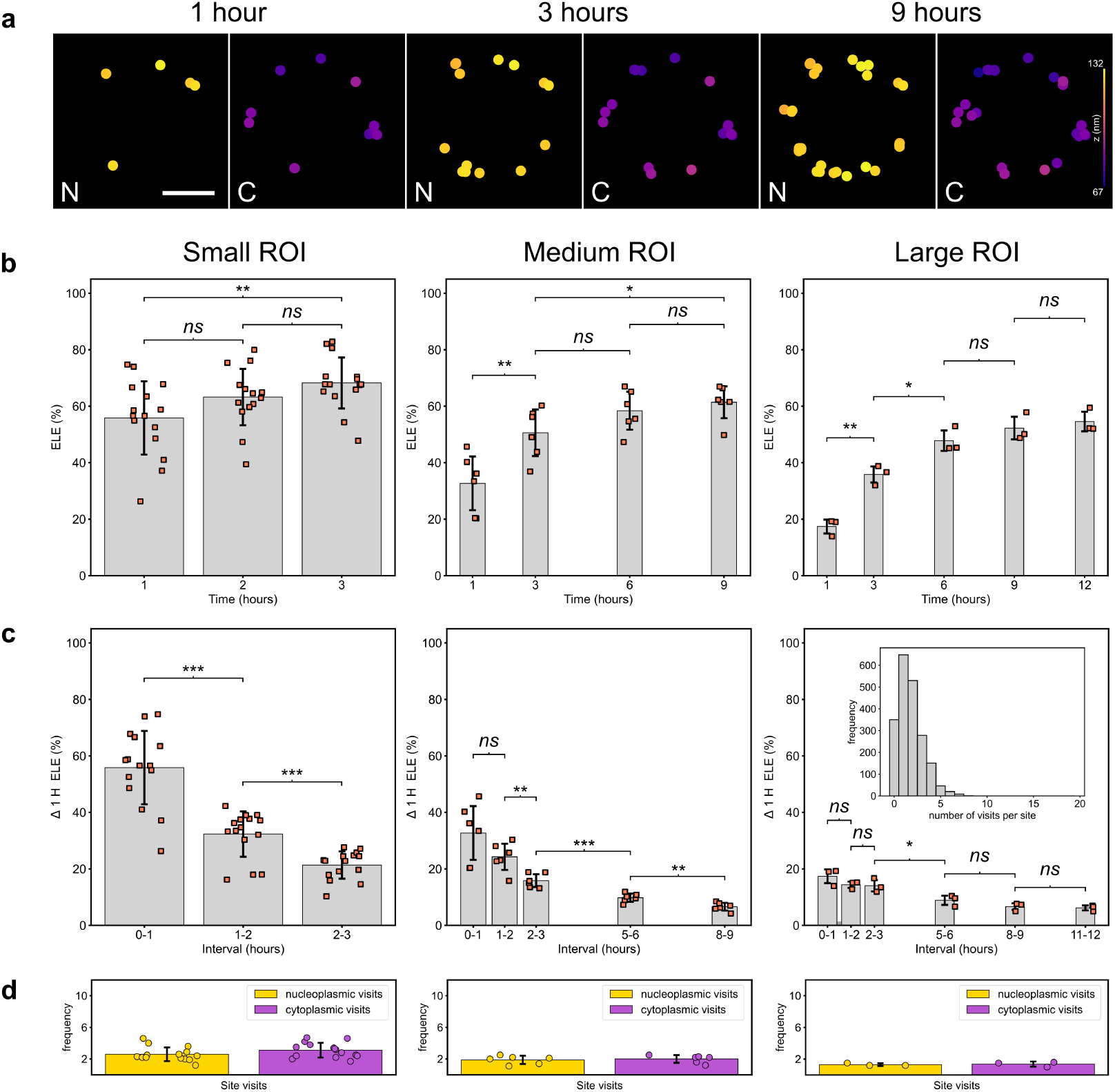
Effective labeling effciency (ELE) over time. a. MINFLUX localizations of Nup96 proteins from nucleoplasmic (N) and cytoplasmic (C) NPC rings at different points in time. As the scan time increases the completeness of the Nup96 distribution for both rings improve with some sites obtaining additional revisits. **b**. Calculated ELE over time for increasing MINFLUX scan-size regions, from small to large, incorporating approximately 10, 50 and 100 NPCs per experimental condition. **c**. ELE measurements taken at one-hour intervals for the same NPCs in **b** demonstrate substantial DNA-PAINT site-loss as the experiment progresses for all scan-sizes. A histogram inset within the data for the ‘large’ ROI demonstrates how the site-loss results in only a small number of revisits to DNA-PAINT marker sites in MINFLUX experiments. **d**. Bar graphs show the number of visits to both nucleoplasmic and cytoplasmic sites for each of the imaging fields of view obtained at their maximal ELE time points: 3, 9, and 12 hours. These data were collected from N independent repeats, n_cell_ imaged cells, and n_NPC_ imaged NPCs. Small ROI: N = 4, n_cell_ = 15, n_NPC_ = 209, medium ROI: N = 4, n_cell_ = 6, n_NPC_ = 280, large ROI: N = 3, n_cell_ = 3, n_NPC_ = 342. Scale bar: 50 nm.

#### ELE with conventional DNA-PAINT

As in other SMLM methodologies, MINFLUX microscopy builds an image gradually over time. In current implementations of MINFLUX microscopy we have the extreme case that one molecule is localized at a time and all localizations are acquired strictly sequentially with no parallel processing (which helps accelerate camera-based widefield SMLM). Accordingly, structures such as the NPC structures imaged here, become gradually more complete as acquisition progresses, as shown in Fig. 3a. As an additional consequence of MINFLUX being a single point scanning system, the size of the region selected for imaging has a direct impact on the time to acquire a “complete” image. We therefore measured the *ELE* of U-2 OS-Nup96-mEGFP NPCs at various time points across several scan region sizes we term “small”, “medium” and “large”, corresponding to ROI areas of approximately 2, 9 and 25 µm^2^ and encompassing around 10, 50, or 100 visible NPCs, respectively. Data was generally acquired for extended time periods, over several hours, and we analyzed subsets of the data with increasing analysis cutoff time *T*_*max*_ by filtering only for events with a time of localization t ≤ *T*_*max*_ in the PYME analysis environment to determine how ELE increases progressively with time.

Within the first hour, using the smallest ROIs, an ELE of 55.9 ± 13.0% was obtained, compared to 32.7 ± 9.5% for the medium size ROI, and 17.4 ± 2.5% for the largest field of view imaged, Figure 3b (all values are reported as mean ± standard deviation for all NPCs measured, see also Supp. Table 1). With MINFLUX DNA-PAINT, the small ROI achieved the highest level of *ELE* after ∼3 hours, 68.2 ± 9.0%, whereas the medium and large ROIs peaked slightly lower with means of 61.4 ± 5.7% and 54.6 ± 3.5% obtained within 9 and 12 hours of recording, respectively (although changes in ELE were not significant for the large ROIs beyond 6h). If experiments are continued beyond these durations, ELE does not increase statistically any further (see, for example the large ROI results in Fig. 3b). This is due to two effects, (1) revisits of the same sites do not increase ELE further and (2) although DNA-PAINT overcomes photobleaching of imagers it does still experience a lowering of localization rates due to site-loss.

**Table 1.**
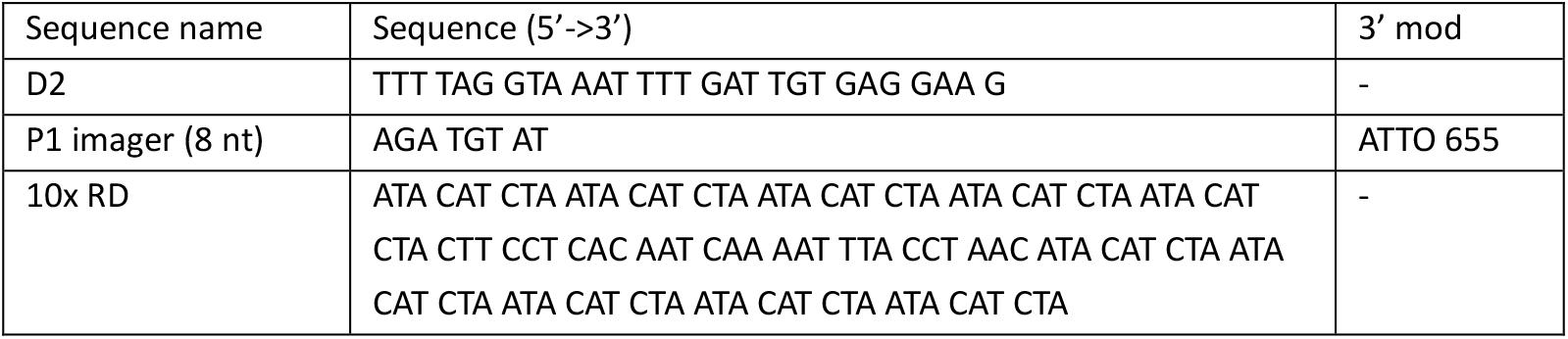
Sequences used for repeat DNA-PAINT.

This effect of site-loss becomes clear by examining how *ELE* was affected by restricting the data used for ELE analysis at hourly intervals as the experiments progressed, in each case only using data from one hour, at analysis starting times of 1, 2 and 3 hours since MINFLUX acquisition began (and several later times with medium and large ROIs), Supp. Table 2. With all ROI sizes we observed a progressive decrease in the *ELE* as compared to measurements taken within the first hour, Figure 3c, which reflects progressive site-loss as also seen by the gradual reduction in event rates (compare also Fig. 1c).

Site-loss has an important effect by limiting the number of revisits of individual target sites. We investigated this by measuring the average number of MINFLUX localizations within a single segment of an NPC that contains two Nup96 target sites (see also schematic in Fig. 1biv,v). Site visit histograms are typically near-exponential in shape, with many sites only visited a single time, especially for medium and large sized ROIs, as illustrated in the inset of Fig. 3c. On average, after an acquisition time at which ELE becomes maximal (3, 6, or 9h for small, medium and large ROIs, respectively) 2.6/3.1± 0.9/0.9 (nucleoplasmic /cytoplasmic side, small ROI), 1.9/2.0 ± 0.5/0.5 (N/C, medium ROI) and 1.3/1.4 ± 0.2/0.3 (N/C, large ROI) visits per site were achieved, as shown in Fig. 3d. Importantly, these numbers do not appreciably increase by longer imaging due to irreversible site-loss (see, for example, Supp Fig. 4).

**Figure 4.**
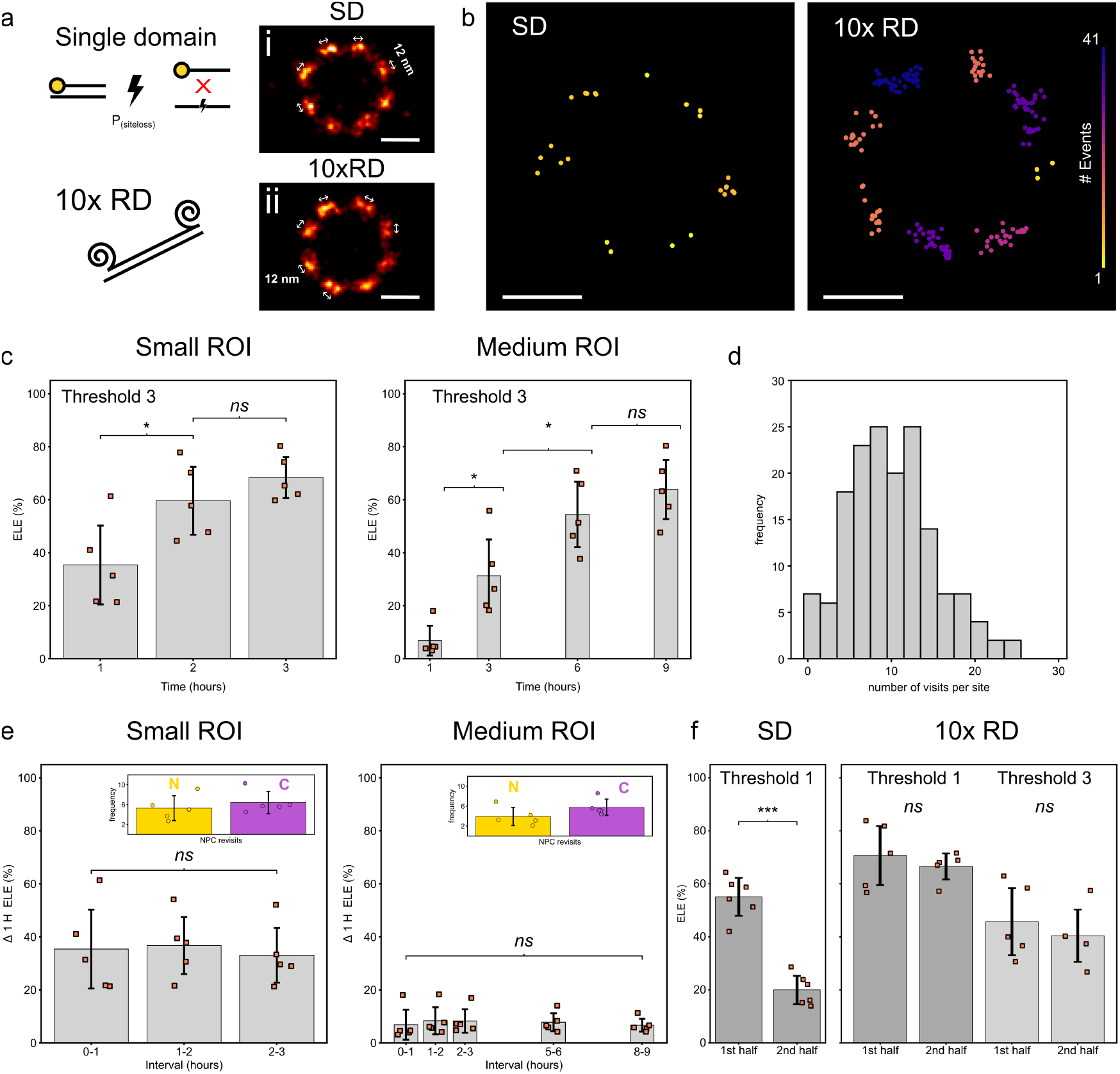
Repeat domain (RD) ELE. a. A schematic depicting the susceptibility for DNA-PAINT docking strands to become photo-damaged during a MINFLUX acquisition. An oligonucleotide consisting of repeat docking sequences fixed to target DNA-PAINT markers provides additional redundancy, increasing the probability of detection as an experiment progresses. Gallery rendered images for **i)** single domain (SD) and **ii)** 10x RD averaged NPC images using template aligned NPCs from two exemplary small ROI datasets. **b**. The additional docking locations results in a greater number of revisits to Nup96 sites, approximately 3-fold. Example localizations from cytosolic NPC rings exhibit a greater number of returns for similar scan sizes at the same time point. **c**. ELE for 10x RD for both small and medium scan sizes using an increased threshold requiring a minimum of three visits per site. **d**. A typical histogram of site visits in this configuration no longer exhibits the near-exponential behavior observed with single-domain site-loss. **e**. ELE measurements taken at one-hour intervals for the same NPCs in **c** demonstrate the effect of having additional redundancy to reduce the observable effects of DNA-PAINT site-loss during MINFLUX imaging. This additionally assists the number of revisits each site obtains, see inset bar graphs for nucleoplasmic (N) and cytoplasmic (C) visits at 3 and 9 hours for the ‘small’ and ‘medium’ ROIs. **f**. ELE measurements for the first and second half of experiments examining a ‘medium’ ROI for normal DNA-PAINT harboring a single domain (SD) demonstrate a significant reduction in the obtainable ELE. Whereas RD DNA-PAINT demonstrates a considerable resistance to site-loss for the same experimental conditions. These data were collected from N independent repeats, n_cell_ imaged cells, and n_NPC_ imaged NPCs. Small ROI: N = 3, n_cell_ = 5, n_NPC_ = 61, medium ROI: N =3, n_cell_ = 5, n_NPC_ = 177, and for the SD medium ROI: N = 4, n_cell_ = 6, n_NPC_ = 280. Scale bars: 50 nm.

Finally, we briefly compared these results with some data from MINFLUX dSTORM experiments. In contrast to MINFLUX DNA-PAINT, in photo-switching MINFLUX dSTORM acquisitions with Alexa Fluor 647, a much smaller *ELE* at the end of experiments (dictated by the near complete decay of localization rates) of 29.5 ± 2.3% was obtained (for medium-sized ROIs, Supp. Figure 5a, see also Supp. Table 3). This coincided with a rapid decrease in the number of localizations detected as the scan proceeded, Supp. Figure 5b. Note that the decay of localization rates can be greatly influenced by the makeup and performance of the inherently labile switching buffers, therefore precise dSTORM results may differ across laboratories.

#### ELE with Repeat DNA-PAINT

Our analysis of MINFLUX data of NPCs with conventional DNA-PAINT docking strands suggests that site-loss may preclude further increasing the detection probability *P*_*detect*_. In addition, from a spatial resolution point of view and to increase confidence in imaging results in SMLM, it is generally desirable to increase the number of site visits at a target site. A promising approach in this context is repeat DNA-PAINT, i.e. use of DNA-PAINT markers with multiple repeat docking domains^12,25,26^, shown in the schematic in Fig. 4a, as established by us and others. Composite rendered images constructed by averaging multiple NPCs for both single domain (SD, Fig 4ai) and repeat domain (RD, Fig 4aii) exhibit clustering suggestive of resolving the 12 nm spacing between individual sites. Markers bearing repeat domains have a number of advantages, including the ability to reduce imager concentrations and accelerate acquisition. Here, we decided to utilize a 10x RD (repeat domain) as we had previously shown that this configuration can greatly reduce the rate of site-loss^12^. In comparison to a marker exhibiting a single domain docking strand, the repeat strands were operated at an approximate order of magnitude reduced imager concentration, which also helps reduce background fluorescence from imagers in solution. When imaged for an extended period of time, ∼9 hours, the repeat domain markers achieved ∼3 times as many visits per site for the equivalent MINFLUX imaging duration, Figure 4b. Despite this, we interestingly found Fourier Shell Correlation (FSC) measurements for SD and RD to be similar to one another at around 9 nm, Supp. Fig. 6. When evaluated with an increased threshold of at least 3 events in a segment (enabled by the much higher number of target site visits), the small ROI gave an ELE of 35.4 ± 14.9% within the first hour, by three hours this had matched the single domain lower threshold at 68.4 ± 7.7%. Similar results were observed with a medium-sized ROI, 63.9 ± 11.2% at 9 hours, Figure 4c, see Supp. Table 4. See also Supp. Fig 7 and Supp. Table 5 to compare RD data at the lower threshold.

A typical histogram of site visits in this configuration has a mode at 10 site visits (Fig. 4d) and has lost the near-exponential appearance resulting from substantial site-loss with conventional DNA-PAINT. In agreement with the idea that site-loss is greatly reduced with this strategy, the measurement of ELE at hourly intervals revealed a steady state of Nup96 detections in both small (35.1 ± 2.1%) and medium (7.6 ± 1.2%) ROIs, Fig. 4e, see also Supp. Table 6. The number of site visits increases with repeat DNA-PAINT to 5.3/6.4 ± 2.5/2.2 (N/C, small ROI) and 3.9/5.8 ± 1.8/1.6 (N/C, medium ROI) after 3 and 9 h, respectively (Fig. 4e, insets). Interestingly, with the increased revisits per site, our repeat domain acquisitions had a slightly elevated number of cytosolic event detection compared to those closer to the nucleus, see also Supp. Figure 8.

When datasets were split in half by acquisition time, the ELE achieved using localizations from the first and second part could be compared. Single domain (SD) DNA-PAINT exhibited a stark contrast to 10x RD experiments with a reduction in ELE 55.1 ± 7.2 % to 20.0 ± 5.3 %, p<<0.001, whereas for 10x RD ELE remains statisticaly indistinguishable at 45.7 ± 12.7 % vs 40.4 ± 9.9 %, p=0.528 (Fig. 4f). Note that with a lower threshold of 1, this increases to a 10x RD ELE of 70.7 ± 11.1 vs 66.6 ± 4.9, p=0.521.

## Discussion

As we improve the obtainable resolution of our microscopy approaches, it is essential to consistently benchmark the quality of the data produced. MINFLUX is a powerful technique, but it requires a complex implementation. With the use of the U-2 OS-Nup96-mEGFP modified cell line we have benchmarked the obtainable ELE using DNA-PAINT single-molecule events on a commercial MINFLUX microscope system. The results and open-source software will enable other users to check the quality of their labeling procedures and determine their individual detection capabilities.

The Nup96 protein is an ideal biological target for quantitative assessment of labeling and experimental conditions, as has been previously outlined.^20^ As we show here, in MINFLUX experiments, not only the chemical labeling capabilities but also the detection probability play a critical role in the obtainable data. Our measurements in small, medium and large MINFLUX scan-size regions demonstrated how easily one can under-sample a super-resolution target by simply stopping the experiment too early. Using a known biological structure was critical in assessing this. The data we have acquired to date and the measurements shown here lead us to favor DNA-PAINT over the conventional single-molecule direct photo-switching of dyes like Alexa Fluor 647 to avoid the labile nature of the photo-switching buffers necessary for lasting blink performance.^6^ In our hands, the dSTORM-type MINFLUX experiments were generally brought to an end prematurely as the fluorophores enter states of permanent photobleaching, leading to low ELE values for even moderate scan sizes. dSTORM-based MINFLUX experiments may be able to achieve similar high ELEs with virtually complete marker detection by using optimized buffers and imaging settings. However by comparison, the DNA-PAINT approach should be more easily replicable across laboratories and, in our experience, efficient marker detection can be robustly achieved by moderately experienced users.

Nevertheless, in MINFLUX DNA-PAINT experiments we also observed a drop in the number of detected events as the experiment proceeded, that are most likely attributable to site-loss. This is an effect that we and others have mostly studied in synthetic samples where there is known to be a single docking site at the marker location.^11,12^ Custom DNA-PAINT IgG secondary antibodies with unknown (typically 1-3) quantities of docking strands and amplification resulting from a primary/secondary AB labeling system, provide some redundancies to photo-damage and, in widefield super-resolution setups, have been demonstrated to produce near constant event rates.^27^ We attribute the noticeable site-loss, observed in our ‘normal’, single domain DNA-PAINT docking strand MINFLUX experiments, to the serial nature of the scanning MINFLUX approach, combined with the use of single-domain nanobodies that exhibit only a single docking strand per marker. With this labeling approach, as a DNA-PAINT experiment progresses over time, the likelihood of returning new data points diminishes. This means that it takes longer to reach the maximum obtainable ELE values and in the worst case, where all sites become damaged and no new additional events are possible, the detection probability never reaches 100%, meaning that the upper limit ELE value set by the chemical labeling efficiency is never reached.

To interpret our results and guide the design of MINFLUX experiments we constructed a very simple phenomenological model that, while clearly neglecting much photo-physical detail, recapitulates our main findings. The model is described in the supplementary methods and its main behavior is illustrated in Supp. Fig. 9. The model predicts that the detection probability should increase over time as a saturating exponential with a time constant that is on the order of the effective revisit time constant. With extensive site-loss, the time-course slightly accelerates, a characteristic parameter is the ratio *α* ≈ τ_revisit_/τ_SL_ as shown in Supp. Fig. 9 c. More importantly, this also affects the saturating value of the detection probability, with increasing site-loss it may be well under 100%. In addition, the site visit histograms from the simple model recapitulate the qualitative transition we observe in the experiments, from an exponential-like behavior with 1 or 2 site visits at most to repeated visits with reduced site-loss in repeat DNA-PAINT.

We note that experimentally we always measure ELE that is the product of chemical and detection probabilities, *ELE* = p_chem_p_detect_. In good approximation, p_chem_ should be constant across experiments, so that we can interpret these to show *ELE* = c_*chem*_ p_detect_, with a constant c_*chem*_ that is in the range 0.7-0.85^20^. With this in mind, MINFLUX standard DNA-PAINT appears to be increasingly affected by site-loss with larger ROI size, implying that with the increase of the effective revisit-time due to the larger ROI, the effective site-loss rate increases relatively even faster. This suggests it may be beneficial to acquire several (to increase sampling) smaller ROIs to saturation (1-2h) rather than one very large ROI which may require ∼9 h. This also avoids potential experimental issues during a single long acquisition that may then ruin the one large dataset. In agreement with this general observation, the number of site visits is also larger with small ROIs, which is desirable.

Overall, standard MINFLUX DNA-PAINT eventually experiences substantial site-loss in all cases, and this was only rectified by using repeat DNA-PAINT which enables sustained acquisitions with many site revisits. The mean number can be adjusted by altering the acquisition duration. In addition, repeat DNA-PAINT allows increasing p_detect_ close to 100%, exploiting the achievable limit set by chemical labeling.

Other approaches could be used to reduce site-loss, either in combination with repeat DNA-PAINT or as alternatives. Imager dye stability may be a factor since it is likely that excited dye molecules in radical states are responsible for site-loss.^11^ Therefore, using more photostable dyes or solution additives to reduce triplet state lifetimes may have a similar effect.^28^

It is possible that other properties of the scan or the sample affect site-loss, such as the effective chemical labeling density. These may influence the ratio of effective revisit times and characteristic site-loss times. For this reason, experiments to measure the time course and maximize p_detect_ should be carried out for the biological sample in question. Due to the purely serial nature of current MINFLUX implementations this requires extensive imaging durations, even for moderate ROI sizes.

Our repeat domain sequences provide 10 identical binding locations for the imager to bind. This feature provides redundancy if any one site becomes damaged during a MINFLUX localization or by being in proximity to one. We observed a near-constant event rate, indicated through the measurement of ELEs with hourly intervals, for small and medium scan sizes. The repeat domains additionally enable the user to reduce imager concentrations, thereby reducing the effects of backgrounds whilst maintaining binding kinetics, and is also compatible with a fluorogenic approach.^15^ Aside from repeat domain approaches, there are alternative ways to reduce site-loss. Recently, the use of cyclooctatetraene (COT) modified oligonucleotides or the addition of COT in imaging buffers were demonstrated to also reduce the effects of photo-damage and could be combined with single or repeat domains to further enhance MINFLUX data acquisitions.^29^

The ELE values we report are comparable to previously reported values using other super-resolution approaches and can be used to quantify other MINFLUX, and similar, experimental setups. The ELE determination reported here could additionally be used in co-cultured samples to quantify the ELE of other GFP-tagged targets by fixing and labeling cells together on the same slide.

## Methods

### Sample preparation

#### Oligonucleotides

Tag X2 anti-GFP single domain nanobodies were purchased from Massive Photonics in either their default ‘Docking Site 3’ and ‘Imager 3’ configuration with undisclosed sequences or as a custom conjugation to the D2 sequence. The repeat domain (10x RD P1) and P1 8nt imager with a 3’ ATTO 655 dye modification were ordered from Eurofins and were HPLC purified. See Table 1 for the oligonucleotide sequences used. The D2 design was previously used to functionalize markers for repeat docking domains^12^ and has additionally been used for other applications.^13,16^

#### Cell culture

U-2 OS-Nup96-mEGFP and Nup96-SNAP cells were purchased from Cytion (clone 195, #300174 & #300444). The cells were grown at 37°C in a humidified incubator containing 5% CO2 in McCoys 5A medium (Fisher Scientific) supplemented with 10% fetal bovine serum (FBS) and 1% Antibiotic-Antimycotic solution (Corning, #30-004-CI). For seeding on coverslips, cells were washed with PBS and incubated with Trypsin-EDTA (0.25%) for 3 minutes until detached. Trypsin was neutralized with fresh medium, and a total of 4×10^4^ cells were plated on 1.5H glass coverslips (Menzel-Gläser 22x22 mm) and allowed to grow for 2 to 3 days before fixation. For fixation, culture medium was removed, and cells were incubated for 10 min at room temperature (RT) in PEM (80mM PIPES, 5mM EGTA, 2mM MgCl_2_) supplemented with 4% formaldehyde (FA) (32% ampoules, Electron Microscopy Sciences, #15714) and 2% sucrose.^30^ Cells were washed 3 times with PBS (Sigma, P4417).

#### Cultured cell sample preparation

Coverslips were attached to custom open-top Perspex chambers using a silicon fast-curing resin. Fixed U-2 OS-Nup96-mEGFP cultured cells were then permeabilized with 0.1% Triton X-100 (Sigma, #T9284) in PBS for 10 minutes at room temperature. After an exchange of buffer to PBS, cells were incubated in Massive Photonics antibody incubation solution for up to 1 hour. The samples were subsequently incubated with 1:200 anti-GFP single-domain nanobody (harboring either Docking Site 3 or custom D2 sequences) for 1 hour at room temperature in Massive Photonics incubation solution. Cells were washed 3 times (∼5 minutes each) with DNA-PAINT buffer (PBS supplemented with an additional 500 mM NaCl (Sigma, S7653). Gold nanoparticles (BBI Solutions, EM.GC150/4) were introduced to the sample 1:1 in DNA-PAINT buffer and allowed to settle onto the coverslip. After sufficient levels of attachment (determined by monitoring the sample on the MINFLUX system and dependent on the sample, aiming for >5 gold particles on the glass surface local to desired imaging regions), samples were washed in DNA-PAINT buffer to remove unbound and jittery gold particles. Occasionally, an additional washing step using 10 mM MgCl_2_ in PBS was used to help remove inadequately bound gold particles that remained on top of cells. Imager 3 was introduced to the sample in a concentration of ∼1 nM diluted in DNA-PAINT buffer.

For the repeat domain experiments, the labeling was conducted as above, but with the addition of incubating the sample with approximately 200 nM 10x RD P1 for 10 minutes at room temperature. The samples were washed several times with DNA-PAINT buffer in order to reduce surplus docking sites. Imaging was then conducted with an order of magnitude reduction in imager concentration (∼0.1 nM) in comparison to experiments without the repeat domains.

#### Photo-switching labeling

For pre-fixation, culture medium was removed, and cells were incubated for 30 seconds at RT with fixation solution consisting of 2.4% Formaldehyde solution (Thermo, 28908) in PBS (Gibco, 14040). Cells were washed once with PBS and permeabilized with 0.4% Triton X-100 (Sigma, T9284) in PBS for 3 minutes at RT. After another wash with PBS, cells were fully fixed for 30 minutes at RT with fixation solution, followed by another wash. Samples were then quenched for 5 minutes at RT with 50 mM NH4Cl (Sigma, 213330) in PBS, followed by three additional 5 minute washes with PBS. To avoid unspecific binding of the labeling agent to the sample, a 30 minute incubation at RT with ImageIT Signal Enhancer (Invitrogen, I36933) was performed. Labeling was achieved by incubating the sample for 1 hour at RT with a 1 µM solution of SNAP-Surface Alexa-647 (NEB, S9136S) diluted in PBS supplemented with 0.5% BSA (Sigma, A9647) and 1 mM DTT (Sigma, D9779). A final round of three washes of 5 minutes each at RT with PBS was performed prior to sample mounting.

Gold nanoparticles (BBI Solutions, EM.GC150/4) were introduced to the sample undiluted and allowed to settle onto the coverslip for 5 minutes at RT. After attachment, samples were washed with PBS several times to remove any unbound gold particles.

Freshly prepared GLOX buffer consisting of 50 mM Tris/HCl (Invitrogen, 15568-025), 10 mM NaCl (Sigma, 71380), 10% (w/v) Glucose (Sigma, G5400), 64 µg/mL catalase (Sigma, C1345) and 0.4 mg/mL glucose oxidase (Sigma, G2133) was adjusted to pH 8.0 and supplemented with 15 mM cysteamine (Sigma, 30070). Samples were then mounted on chambered slides (BMS, 12290) filled with approximately 100 µL of imaging buffer and secured with silicone glue (Picodent eco-sil).

### Data acquisition

#### Microscope setup

MINFLUX data were acquired using a commercial 3D MINFLUX microscope (Abberior Instruments) equipped with 405 nm, 485 nm, 561 nm, and 642 nm laser lines. Additionally, a 980 nm IR laser and a PI nano® XYZ Stage were used for active sample stabilization.

Acquisitions were performed using a 100x 1.45 NA UPlanXAPO oil objective (Olympus), and the pinhole was set to 0.83AU.

Fluorophores were excited using the 642 nm laser, and fluorescence was detected *via* two APDs with different spectral windows: Cy5 near (650 - 685 nm), and Cy5 far (685 - 720 nm). The default 3D Imaging sequence provided by Abberior was used to determine the localizations of the fluorescent molecules. This sequence defines the parameters of each iterative localization process, such as the pattern type, the diameter of the targeted coordinate pattern (TCP), and the minimal number of photons required for valid localization, see also Supp. Table 7).

#### PSF check and alignment

Before each acquisition, PSF and pinhole alignment were checked. The red beads from a nanoparticle gold (150 nm) + 2C fluor (120 nm) slide (Abberior #NP-3012) were used to assess the PSF shape. Briefly, 2D and 3D donut-shaped patterns were examined to verify the presence of a zero-intensity center in the x-y, x-z, and y-z planes. Additionally, the donut homogeneity was assessed visually. Pinhole alignment was assessed using red beads, ensuring that the 3D PSF would not be clipped when setting the pinhole to the experimental settings (0.83AU).

### DNA-PAINT MINFLUX experiments

Confocal scans were collected prior to MINFLUX acquisitions to capture the diffraction-limited distribution of NPCs using a 488 excitation laser and STAR488 Quad Band filter cube (spectral range 500nm – 550nm). A qualitative assessment of imager binding kinetics was performed by recording an *xyt* sequence of 50-100 frames for a 3×3 µm region of interest (ROI) with 30 µs dwell time and 30 nm pixel size. These settings enabled clearly visualized circular spots on the scans as imagers were immobilized at the docking sites, see Supp. Media 1. Using the visual display, ROIs were selected for ∼1.5x1.5 µm (small), ∼3x3 µm (medium), or ∼5x5 µm (large) boxed areas for MINFLUX scanning. Depending on the ROI size (small, medium, large) the experiment was conducted for 3, 9 or 12 hours respectively. For all experiments the MINFLUX beamline monitoring (MBM) tracking module was used on selected gold particles. Each experiment aimed to have at least 5 gold particles tracked for the duration of the acquisition. The Abberior Instruments 642 nm laser was nominally operated at a software setting of ∼10% power with a recorded measurement before the periscope of ∼ 5.8mW.

#### Photo-switching MINFLUX experiments

Confocal scans were collected prior to MINFLUX acquisitions to capture the diffraction-limited distribution of NPCs using a 640 excitation laser and the Cy5 near and Cy5 far detectors (total detection window: 650-720 nm). Using the visual display, ∼ 3x3 µm (medium) ROIs were selected for MINFLUX scanning. Prior to the measurement, off-switching of each ROI was performed by scanning with high laser power (software setting of ∼ 10-20%, measured power of ∼ 15-20 µW at the sample) and slow scanning speed (pixel size of 10 nm and dwell time of 5 µs) until only sparse diffraction limited events were observed.

For all experiments the MINFLUX beamline monitoring (MBM) tracking module was used on selected gold particles. Each experiment aimed to have at least 5 gold particles tracked for the duration of the acquisition. The Abberior Instruments 640 nm laser was nominally operated at a software setting of ∼6% power with a recorded measurement of ∼ 7 µW at the sample. 405 nm activation laser power was gradually increased during the acquisition to ensure a consistent rate of detections while maintaining the single-molecule detection regime.

#### Data analysis and visualization

Exported .npy or .zarr format data from the Abberior software ‘Imspector’ were imported into the open-source Python Microscopy Environment (PyME) visualization and analysis software, http://github.com/python-microscopy/python-microscopy, with additional functional plug-ins from https://github.com/csoeller/PYME-extra. Upon import into PYME a foreshortening factor of 0.72 was applied to all z coordinates, similar as done previously^31^. This corrected the separation between cytoplasmic and nucleoplasmic rings from 69.3 ± 7.2 to 50.8 ± 5.8 nm, Supp. Figure 10. MBM tracks were plotted to inspect their agreement with one another and divergent tracks were rejected. To reduce poor localizations and also to reject likely multi-emitter events from overlapping emissions, CFR and EFO filtering were conducted, as described previously. Non-specific labeling of the sample, as previously reported,^12,16^ can still be detected as a true event and was therefore considered for threshold choice in determining NPC labeling, see below. Localizations from the same trace were coalesced to a single detection based on the inherent ‘traceID’ property, as described previously^9^. The full dataset was rendered as a 20 nm pixel size Gaussian rendered image^32^ to identify and create a mask of super-resolved NPCs. This mask was then injected back into the PyME pipeline to obtain events within the mask and each NPC was assigned a unique ‘objectID’ to mark these events.

### Quantitative Determination of ELE by maximum-likelihood fitting of NPC templates

#### NPC mask creation

Using the full duration experiment, drift-corrected localization events were rendered as 2D Gaussian images with 10 nm pixel size and saved as .tif files. These images were imported into the software package Fiji, ImageJ,^33^ using Bioformats.^34^ The circular selection tool was used to semi-automatically select identifiable NPCs (excluding NPCs that are partially clipped at MINFLUX ROI edges), recorded using the “ROI Manager” tool and saved as .zip file. In PyME, the previously rendered image the imported Fiji ROI were used to create a mask of the NPC locations and subsequently created unique object identifiers (IDs) to tag all events associated with NPCs in the mask.

#### NPC analysis

For each NPC the associated set of localizations (identified by the unique ID label) were fit to a 3D double ring template using a maximum-likelihood algorithm following an approach described previously^24^. The 3D template was generated from a geometric arrangement of two rings (nominal diameter 107 nm) spaced 50 nm apart axially. These were convolved with a Gaussian (sigma = 5 nm) in 3D to provide a smooth alignment template. A small, uniformly distributed probability was added to the whole template volume dataset to account for background localizations^24^. We adopted a “smooth” model versus a “detailed” NPC template for robust parameter estimation^24^. Similarly to previous work, also for robustness, we do not attempt to resolve the pairs of nearby Nup96 sites (laterally ∼12 nm apart) in the 8-fold symmetry NPC structure, but rather test for labeling of each of the 8 radial “segments” that each contain these site pairs^20^. Since we have 3D data, we do this separately for cytoplasmic and nucleoplasmic sides of the NPC, i.e. checking for labeling of 8 cytoplasmic and 8 nucleoplasmic segments, allowing for labeling of up to 16 segments in total per NPC.

For each NPC and its associated localizations the negative logarithm of the likelihood (NLL) is minimized, as described^24^, to maximize the likelihood. To minimize the NLL, 7 parameters are varied which describe coordinate transformations to line up the MINFLUX localizations with the template, using a global minimizer (the `basinhopping` function of the `scipy.optimize` package), as described^9^ and as illustrated in Fig. 2b.

#### Segment analysis

To check for segment labeling a threshold of 1 or 3 compound localizations was used to regard a segment as labeled (see threshold determination below). Prior to segment labeling analysis the aligned NPC data was additionally rotationally aligned to “segment boundaries” as described.^9^

#### Effective labeling efficiency

To determine the ELE value for a MINFLUX 3D dataset the template fitting procedure and segment labeling testing were conducted for each NPC in the dataset. The data were then pooled to generate a histogram of the number of segments that were labeled across the dataset, as shown in Fig. 2e, displaying the relative frequency of 0 to 16 segments being detected as labeled. Finally, for obtaining the ELE value the probabilistic model described in the supplementary model was fit to the data histogram in cumulative form, as shown in Fig. 2f. The best fit parameter *P*_*LE*_ of the model, together with its uncertainty *ΔP*_*LE*_, are then provided as an estimate of ELE for the given dataset.

The complete NPC fitting and ELE estimation procedure is implemented in the `PYMEcs.Analysis.NPC` module of the `PYME-extra` package. A graphical interface is provided by the `NPCcalcLM` module, also part of the `PYME-extra` package, as a plugin for PYMEVisualize.

#### Gallery mode, averaging NPCs

To produce an “average” image or a gallery of NPCs from a dataset, all normalized coordinates obtained from fitting to the NPC template and rotational segment alignment were overlaid in PYMEVisualize (using the ‘Add NPC Gallery’ command) where (a) it could be chosen to overlay all events in a single coordinate system at the origin or (b) NPCs and/or nucleoplasmic/cytoplasmic segments were arranged next to each other on a regular grid.

Functionality (a) was used to generate an average image of all aligned NPC data by rendering all overlaid NPC localizations using Gaussian rendering (pixel size 2 nm, Gaussian standard deviation 2 nm, e.g. images in Fig. 4b). Functionality (b) was used to generate a gallery of NPCs next to each other for visual inspection with segment boundaries overlaid (e.g. the NPC sample shown in Fig. 2e, displaying cytoplasmic and nucleoplasmic localizations side by side for visual inspection).

#### Fourier Shell Correlation Measurements

The 3D MINFLUX data was split into two blocks using the “time blocking” functionality of PYMEVisualize and rendered as separate Gaussian-generated volume images using a Gaussian width of 2 nm with a voxel size of 2 or 3 nm. These were then saved as MRC format files using functionality in PYME-extra (“Save MRC volumes for FSC”) and uploaded to the online Electron Microscopy Data Bank FSC server (https://www.ebi.ac.uk/emdb/validation/fsc/, accessed 14/05/25). The downloaded .xml result files were then processed for plotting using a Jupyter Python notebook.

#### Background and threshold determination

Background localizations above and below the NPC focal plane were used to determine the background contribution to our ELE analysis and assist in setting the threshold value.

Background localizations from single domain experiments were analyzed with the same NPC mask used for the specific dataset, but with the NPC localizations removed and only using the background localizations. The change in ELE (ΔELE) was determined by purposely shifting the background events into the NPC localization domain by shifting their z-coordinates into the NPC region and subsequently subtracting the obtained ELE value with these additional background localizations from the previously obtained ELE values without these additional background events. For standard DNA-PAINT markers these values were evaluated for a threshold of 1 compound event and indicated very small contributions from background localizations Supp. Fig. 3a,b. The repeat DNA-PAINT experiments used an ∼order of magnitude reduced imager concentration leading to a reduced background making estimates for background-only measurements difficult. The background localizations from relatively homogeneous datasets were taken from above and below the NPCs and their coordinates were axially transformed into the NPC signal. The ΔELE was then measured as described above using a threshold of 1 compound localization and 3 compound localizations per segment, Supp. Fig. 3c,d.

#### Further analysis

Custom Jupyter (https://jupyter.org) notebooks written in python 3.7.13 code were used to create the plots throughout this manuscript and supporting information, see data availability comment.

#### Statistics and reproducibility

Data is given as the mean values +/-standard deviation. All p-values were obtained using the Python library package SciPy (version 1.3.0) and the ‘stats’ module by running independent t-tests using the function ‘ttest_ind’. A p-value <0.05 was considered as statistically significant and represented on figures as *, p<0.01 as **, and p<0.001 as ***, *ns* denoted non-significance.

## Acknowledgements

The authors would like to acknowledge funding support from the Swiss National Science Foundation (SNSF 310030_208109 to C. S.). The equipment was supported by the SNSF (SNSF R’Equip 316030_213543 to C. S.), the University of Bern Innovation Fund and the Faculty of Medicine. We thank Evelina Lučinskaitė and Gabriele Bleuer for assistance in culturing cells.

## Author contributions

Conceptualization was done by C.S and A.H.C. Data curation was performed by C.S and

A.H.C. Formal analysis was done by C.S, A.F.E.B and A.H.C. Funds were acquired by C.S. Investigation was performed by C.S, A.F.E.B, J.C and A.H.C. Methodology was developed by C.S, A.F.E.B and A.H.C. Project administration was done by C.S and A.H.C. Resources were acquired by C.S, A.F.E.B, J.C and A.H.C. Software handling was done by C.S and A.H.C. Supervision was performed by C.S and A.H.C. Validation was done by C.S, A.F.E.B and A.H.C. Visualization was performed by C.S and A.H.C. All authors contributed to the writing of the manuscript.

## References

1. Balzarotti, F. et al. Nanometer resolution imaging and tracking of fluorescent molecules with minimal photon fluxes. Science 355, 606–612 (2017).

2. Hell, S. W. & Wichmann, J. Breaking the diffraction resolution limit by stimulated emission: stimulated-emission-depletion fluorescence microscopy. Opt. Lett. 19, 780– 782 (1994).

3. Klar, T. A., Jakobs, S., Dyba, M., Egner, A. & Hell, S. W. Fluorescence microscopy with diffraction resolution barrier broken by stimulated emission. Proc. Natl. Acad. Sci. 97, 8206–8210 (2000).

4. Rust, M. J., Bates, M. & Zhuang, X. Sub-diffraction-limit imaging by stochastic optical reconstruction microscopy (STORM). Nat. Methods 3, 793–796 (2006).

5. Betzig, E. et al. Imaging Intracellular Fluorescent Proteins at Nanometer Resolution. Science 313, 1642–1645 (2006).

6. Dempsey, G. T., Vaughan, J. C., Chen, K. H., Bates, M. & Zhuang, X. Evaluation of fluorophores for optimal performance in localization-based super-resolution imaging. Nat. Methods 8, 1027–1036 (2011).

7. Jungmann, R. et al. Single-Molecule Kinetics and Super-Resolution Microscopy by Fluorescence Imaging of Transient Binding on DNA Origami. Nano Lett. 10, 4756–4761 (2010).

8. Ostersehlt, L. M. et al. DNA-PAINT MINFLUX nanoscopy. Nat. Methods 19, 1072–1075 (2022).

9. Clowsley, A. H. et al. Analysis of RyR2 distribution in HEK293 cells and mouse cardiac myocytes using 3D MINFLUX microscopy. 2023.07.26.550636 Preprint at 10.1101/2023.07.26.550636 (2023).

10. Jungmann, R. et al. Multiplexed 3D cellular super-resolution imaging with DNA-PAINT and Exchange-PAINT. Nat. Methods 11, 313–318 (2014).

11. Blumhardt, P. et al. Photo-Induced Depletion of Binding Sites in DNA-PAINT Microscopy. Mol. Basel Switz. 23, 3165 (2018).

12. Clowsley, A. H. et al. Repeat DNA-PAINT suppresses background and non-specific signals in optical nanoscopy. Nat. Commun. 12, 501 (2021).

13. Clowsley, A. H. et al. Detecting Nanoscale Distribution of Protein Pairs by Proximity-Dependent Super-resolution Microscopy. J. Am. Chem. Soc. 142, 12069–12078 (2020).

14. Brockman, J. M. et al. Live-cell super-resolved PAINT imaging of piconewton cellular traction forces. Nat. Methods 17, 1018–1024 (2020).

15. Chung, K. K. et al. Fluorogenic DNA-PAINT for faster, low-background super-resolution imaging. Nat. Methods 19, 554–559 (2022).

16. Lučinskaitė, E. et al. Reduced Non-Specific Binding of Super-Resolution DNA-PAINT Markers by Shielded DNA-PAINT Labeling Protocols. Small 20, 2405032 (2024).

17. Prakash, K. At the molecular resolution with MINFLUX? Philos. Trans. R. Soc. Math. Phys. Eng. Sci. 380, 20200145 (2022).

18. Prakash, K. & Curd, A. P. Assessment of 3D MINFLUX data for quantitative structural biology in cells. Nat. Methods 20, 48–51 (2023).

19. Gwosch, K. C. et al. Reply to: Assessment of 3D MINFLUX data for quantitative structural biology in cells. Nat. Methods 20, 52–54 (2023).

20. Thevathasan, J. V. et al. Nuclear pores as versatile reference standards for quantitative superresolution microscopy. Nat. Methods 16, 1045–1053 (2019).

21. Marin, Z. & Ries, J. Evaluating MINFLUX experimental performance in silico. 2025.04.08.647786 Preprint at 10.1101/2025.04.08.647786 (2025).

22. Schindelin, J. et al. Fiji: an open-source platform for biological-image analysis. Nat. Methods 9, 676–682 (2012).

23. Marin, Z. et al. PYMEVisualize: an open-source tool for exploring 3D super-resolution data. Nat. Methods 18, 582–584 (2021).

24. Wu, Y.-L. et al. Maximum-likelihood model fitting for quantitative analysis of SMLM data. Nat. Methods 1–10 (2022) doi:10.1038/s41592-022-01676-z.

25. Wade, O. K. et al. 124-Color Super-resolution Imaging by Engineering DNA-PAINT Blinking Kinetics. Nano Lett. 19, 2641–2646 (2019).

26. Schueder, F. et al. An order of magnitude faster DNA-PAINT imaging by optimized sequence design and buffer conditions. Nat. Methods 16, 1101–1104 (2019).

27. Lutz, T. et al. Versatile multiplexed super-resolution imaging of nanostructures by Quencher-Exchange-PAINT. Nano Res. 11, 6141–6154 (2018).

28. Steen, P. R. et al. The DNA-PAINT palette: a comprehensive performance analysis of fluorescent dyes. Nat. Methods 21, 1755–1762 (2024).

29. Scheckenbach, M. et al. Minimally Invasive DNA-Mediated Photostabilization for Extended Single-Molecule and Super-resolution Imaging. 2025.01.08.631860 Preprint at 10.1101/2025.01.08.631860 (2025).

30. Jimenez, A., Friedl, K. & Leterrier, C. About samples, giving examples: Optimized Single Molecule Localization Microscopy. Methods 174, 100–114 (2020).

31. Gwosch, K. C. et al. MINFLUX nanoscopy delivers 3D multicolor nanometer resolution in cells. Nat. Methods 17, 217–224 (2020).

32. Baddeley, D., Cannell, M. B. & Soeller, C. Visualization of localization microscopy data. Microsc. Microanal. 16, 64–72 (2010).

33. Schindelin, J. et al. Fiji: an open-source platform for biological-image analysis. Nat. Methods 9, 676–682 (2012).

34. Linkert, M. et al. Metadata matters: access to image data in the real world. J. Cell Biol. 189, 777–782 (2010).

