## Supplemental Information for "Optimizing effective labeling efficiency in MINFLUX 3D DNA-PAINT microscopy by maximizing marker detection probability"

### Supplementary Methods

#### A simple model for effective labeling efficiency in DNA-PAINT

Below, we describe a deliberately simplistic and partly phenomenological model with many details removed (as we generally do not know the photo-physical detail underlying some of the processes) to describe the detection component of the effective labeling efficiency in MINFLUX DNA-PAINT imaging. As a starting point, we consider $P_{ELE}$ as resulting from the probability of chemically labeling a site/target, $P_{chem}$, and the probability $P_{detect}$ of detecting valid MINFLUX events from such a marker, which are largely independent, therefore

$$P_{ELE} = \mathrm{P_{chem}P_{detect}}$$

$\mathrm{P_{chem}}$ is a number that typically depends on the affinity of the marker system used and its ability to access the site/target. Based on previous measurements with a similar anti-GFP sdAB marker system in wide-field super-resolution microscopy (where it should be possible to achieve $\mathrm{P_{detect}}$ approaching 100%) we expect it to be around 0.7-0.85[1].

$P_{detect}$ is a function of the total acquisition time T and we expect it to saturate for large $T$, as either (1) previously visited target sites are just revisited or (2) labeling events cease due to a process like site-loss.

We focus on $\mathrm{P_{detect}(T)}$ which we just call $\mathrm{P(T)}$ below. Depending on site-loss and detection stringency (i.e. the threshold number of events), the saturation value may be less than 1. The goal in MINFLUX DNA-PAINT imaging of such marker distributions is to control acquisition so that $P(T)$ gets as close to 1 as possible within a reasonable total time $T$.

##### *Assumptions of the simplified model*

We assume that all key processes can be captured with single time constants for the specific process. Specifically, we use a mean imager attachment time $\tau_{\mathrm{attach}}$, a time between visits $\tau_{\mathrm{revisit}}$ and a site-loss time constant $\tau_{\mathrm{SL}}$.

For widefield DNA-PAINT, $\tau_{\mathrm{attach}} = 1/\mathrm{k_{off}}$, $\tau_{\mathrm{revisit}} = 1/(\mathrm{k_{on}} * \mathrm{imager\_conc})$, i.e. these are directly related to the hybridization on- and off-rates, and individual attachment and revisit times are known to be distributed exponentially around these means[2]. For MINFLUX DNA-PAINT, we make the simplifying assumption that these processes can still be described by equivalent effective time constants, but now the revisit time constant in particular depends in some way on the ROI size and possibly other variables of the MINFLUX scan, i.e. $\tau_{\mathrm{revisit,MINFLUX}} = \mathrm{f(ROI, \ldots)} * \tau_{\mathrm{revisit,DNA-PAINT}}$. Here, $\mathrm{f(ROI, \ldots)}$ is some possibly large factor since in a MINFLUX scan we are spending only a small fraction of the time looking for attachments at a specific location.

$\tau_{SL}$ may depend on the scan ROI, possibly the density of markers in the vicinity of a site, the size of the doughnut, possibly also the concentration of imagers, and the excitation intensity. The main point of the simplified model is that we can describe the number of available sites approximately as $\exp(-T/\tau_{SL})$ where T is the time since the scan was started.

We also neglect backgrounds in this basic model, which seems warranted as the number of background localizations in 3D MINFLUX DNA-PAINT data seems very low.

##### *Describing P(T) as a function of the 3 time constants*

We assume a minimum attachment time $t_{attach,min}$ that is required for detection, typically this would be a fraction of the mean attachment time $\tau_{attach}$, e.g. a third or a tenth, corresponding to, say, 10-50 ms for typical DNA-PAINT imager attachment times; imager attachments shorter than $t_{attach,min}$ just go undetected (in MINFLUX terminology the localization sequence iterations do not complete or the attachment is not even recognized to start an iteration sequence).

In this scenario, we make the estimate that, without site-loss, the probability $P(T)$ of having observed a detected event at a labeled site after some time $T$, is approximately the probability that the revisit time is $t_{revisit} \leq T - t_{attach,min}$, i.e. that there is enough time $t_{attach,min}$ "left over" for an opening to occur. This, for the exponentially distributed revisit time with mean $\tau_{revisit}$ is

$$P(T) = P(t_{revisit} \leq T - t_{attach,min}) =$$

$$1 - \exp(-(T - t_{attach,min})/\tau_{revisit}) = 1 - \exp(T/\tau_{revisit}) * \exp(t_{attach,min}/\tau_{revisit})$$

which is in good approximation

$$P(T) = 1 - \exp(-T/\tau_{revisit})$$

since $t_{attach,min}/\tau_{revisit}$ is very small. We also note that the probability density of this cumulative probability is

$$pdf(t) = 1/\tau_{revisit} * \exp(-t/\tau_{revisit})$$

A simple Monte Carlo model simulating 1000 sites with exponentially distributed attachment and revisit times (see also below) simulates this numerically using a specific $t_{attach,min}$ and the net outcome is that the actual time course of $P(T)$ is well approximated by $1 - \exp(-T/\tau_{eff})$ with an "effective" time constant $\tau_{eff}$ a little larger than $\tau_{revisit}$ for typical detectable minimum imager attachment times, e.g.$\tau_{eff} = 1.25 \times \tau_{revisit}$. If $t_{attach,min}$ is a large multiple of $\tau_{attach}$, then this can lead to substantial lengthening of the time constant as compared to $\tau_{revisit}$ but is not a case we would expect in practice with DNA-PAINT as we design mean attachment times to be well sampled by the type of microscopy we employ (here MINFLUX). We accordingly use

$$P(T)_{no\_SL} = 1 - \exp(-T/\tau_{eff})$$

with the probability density

$$pdf(t) = 1/\tau_{eff} * \exp(-t/\tau_{eff})$$

for all further calculations below.

We extend this to the case with site-loss by assuming that the fraction of remaining available sites after a time $t$ since the scan began is described as $\mathrm{r(t)} = \exp(-\mathrm{t}/\tau_{\mathrm{SL}})$. Using this expression, we obtain the probability to have detected an event with site-loss after an acquisition time T as $\int_0^{\mathrm{T}} \mathrm{r(t)} * \mathrm{pdf(t)dt}$. This works out as

$$\mathrm{P(T)_{SL}} = \mathrm{f_{max}} * \left(1 - \exp(-\mathrm{T}/\tau'_{\mathrm{eff}})\right)$$

with $\mathrm{f_{max}} \leq 1$ and $\tau'_{\mathrm{eff}} < \tau_{\mathrm{eff}}$, i.e. a new effective slightly "faster" $\tau'_{\mathrm{eff}}$, both are functions of the characteristic dimensionless parameter $\alpha = \tau_{\mathrm{eff}}/\tau_{\mathrm{SL}} \ \approx \tau_{\mathrm{revisit}}/\tau_{\mathrm{SL}}$:

$$\mathrm{f_{max}} = \frac{1}{1 + \tau_{\mathrm{eff}}/\tau_{\mathrm{SL}}}$$

$$\tau'_{\mathrm{eff}} = \frac{\tau_{\mathrm{eff}}}{1 + \tau_{\mathrm{eff}}/\tau_{\mathrm{SL}}}$$

which, as expected, reduces to $\mathrm{P(T)_{no\_SL}}$ for $\tau_{\mathrm{SL}} \to \infty$. $\mathrm{f_{max}}$ limits the maximal attainable detection efficiency. The ratio of $\alpha = \tau_{\mathrm{eff}}/\tau_{\mathrm{SL}}$ determines the type of behavior we see:

- $\alpha \lesssim 0.05$ (i.e. the site-loss time course is much slower than the revisit time): repeat DNA-PAINT-like behavior, we can get close to $\mathrm{P_{detect}} = 1$, many detections per site with long acquisitions are possible

- $\alpha \approx 0.1..0.3$ : standard DNA-PAINT-like behavior, $\mathrm{P_{detect}}$ can get close to 1 with long enough acquisition, not too many detections per site

- $\alpha \geq 0.5$ (i.e. the site-loss time course is approaching the revisit time): broadly speaking, dSTORM-like behavior, $\mathrm{P_{detect}}$ remains well below 1, acquisition naturally ceases after a certain time, very few detections per site

##### *Model limitations*

The simplistic model developed above and the Monte Carlo simulations below strictly speaking only apply to widefield DNA-PAINT, where exponentially distributed attachment and revisit times are well-established. For MINFLUX DNA-PAINT, we suggest that revisit times are still approximately exponentially distributed at any given docking site, although the "effective" mean revisit time is (possibly greatly) lengthened by the MINFLUX ROI size and other scan variables/properties. The actual site-loss time course may also deviate somewhat from a simply mono-exponential decay, but for our basic description it should suffice that it captures a monotonic decay with a somewhat characteristic time constant $\tau_{\mathrm{SL}}$. Similarly, when suggesting that $\alpha \geq 0.5$ describes "dSTORM MINFLUX behavior", this is meant in a purely qualitative sense (i.e. generating a scenario where permanent photobleaching is relatively fast as compared to the time to build up structures during acquisition), as details of "blink-on" times (corresponding loosely to imager attachment times) and "re-blink" times (loosely corresponding to revisit times in DNA-PAINT) depend on the photophysics of photoswitching in possibly very different ways from that of DNA-PAINT revisits and site-loss.

Given these limitations, the primary usefulness of the model is (a) in the use of relatively simply interpretable time constants, (b) that we can identify these time constants with the time-course over which $\mathrm{P_{detect}(T)}$ saturates and (c) revealing that the ratio of $\alpha = \tau_{\mathrm{eff}}/\tau_{\mathrm{SL}} \ \approx \tau_{\mathrm{revisit}}/\tau_{\mathrm{SL}}$ is useful to predict the qualitative behavior of the progressing MINFLUX DNA-PAINT acquisition and how this may limit the attainable detection probability $P_{detect}$.

*Monte Carlo simulations to numerically confirm the simple model behavior*

The basic behavior of the simplistic model was implemented as Monte Carlo simulation that generates temporal trajectories for uniform sites with exponentially distributed attachment and revisit times given time constants $\tau_{attach}$, $\tau_{revisit}$ and a minimum detection time $t_{attach,min}$. In addition, it chooses for each temporal trajectory a site-loss time that is exponentially distributed with mean $\tau_{SL}$. Beyond this time, the site becomes "silent" and can no longer experience imager attachment. A few sample trajectories are shown in Fig. SI 9 a. 1000 such trajectories were generated each for a number of parameter settings reflecting different values of $\alpha = \tau_{eff}/\tau_{SL} \approx \tau_{revisit}/\tau_{SL}$, Fig. SI 9 b. The simulated data was evaluated to determine $P(T)$ for thresholds of 1 (Fig. SI 9 c) and 3 visits at minimum (Fig. SI 9 d), and the average number of visits per site was determined, Fig. SI 9 e. The analysis confirms that (1) P(T), for a threshold number of 1 visit per site, is well approximated by an exponential rise with a slightly slowed $\tau_{eff}$ that is a multiple of $\tau_{revisit}$ by a factor only slightly above 1, (2) that an increasing rate of site-loss leads to a slight abbreviation of the time constant with which the asymptotic value is approached, and that (3) this increasing site-loss leads to a reduction in the maximally attainable P(T), as described by the factor $f_{max}$ introduced above; there is also a "foot" indicating a delay before P(T) starts to increase; (4) increasing detection stringency (threshold = 3) leads to a slowing of the time course, as expected (since more "revisit" times are needed), and further reduces the maximally attainable P(T). Finally, (5) the progression of the site-visit histogram from a shape with a clear mode to an exponential-like decay with very few revisits (if any) with increasing alpha is very apparent. In other words, the more comparable the site-loss time becomes to the average revisit time, the fewer revisits are observed, even for extended acquisition times.

#### Probabilistic model for Nup96 labeling of NPC structures

The probabilistic model described below was used to predict the fractions of 0 to 16 labeled segments as a function of the parameter $p_{LE}$ that quantifies the labeling efficiency. The model is obtained following an approach described previously[1] but adapted to the MINFLUX 3D scenario with 8 "segments" each on the cytoplasmic and nucleoplasmic sides of the NPC, respectively, i.e. 16 in total. Starting from the binomial probability distribution of having exactly *k* successes when attempting to label *n* independent sites with a labeling probability $p_{LE}$, which we capture as a function $f(k, n, p_{LE})$,

$$f(k, n, p_{LE}) = \binom{n}{k} p_{LE}^{k} (1 - p_{LE})^{n-k}.$$

Similarly to previous work, we test for labeling of each segment in the 8-fold symmetric NPC structure, with each segment containing two nearby Nup96 sites (~12 nm lateral distance)[1]. The probability $p_{seg,unlabeled}$ that a segment of the NPC structure is unlabeled is obtained from the binomial probability that both Nup96 sites are not labeled

$$p_{seg,unlabeled} = f(0,2, p_{LE})$$

with the probability $p_{seg,labeled}$ that at least one of the sites is labeled as

$$p_{seg,labeled} = 1 - p_{seg,unlabeled} = 1 - f(0,2, p_{LE}).$$

From this expression one obtains the probability $p(N, p_{LE})$ that *N* out of 16 segments are labeled as

$$p(N, p_{LE}) = f(N, 16, p_{seg,labeled}) = f\big(N, 16, 1 - f(0,2, p_{LE})\big).$$

The cumulative probability $p_c(N, p_{LE})$ which is generally shown in plots and used as a fitting function, is then

$$p_c(N, p_{LE}) = \sum_{k=0}^{N} p(k, p_{LE})$$

We process both experimental and predicted probability histograms in cumulative form since the cumulative curves have a characteristic sigmoidal shape that shifts to the right with increasing labeling efficiency. This aids visual inspection of effective labeling performance and fitting. Finally, the best-fit parameter $p_{LE}$ together with its uncertainty $\Delta p_{LE}$ are provided as an estimate of effective labeling efficiency.

#### References


1. Thevathasan, J. V. *et al.* Nuclear pores as versatile reference standards for quantitative superresolution microscopy. *Nat. Methods* **16**, 1045–1053 (2019).

2. Jungmann, R. *et al.* Multiplexed 3D cellular super-resolution imaging with DNA-PAINT and Exchange-PAINT. *Nat. Methods* **11**, 313–318 (2014).

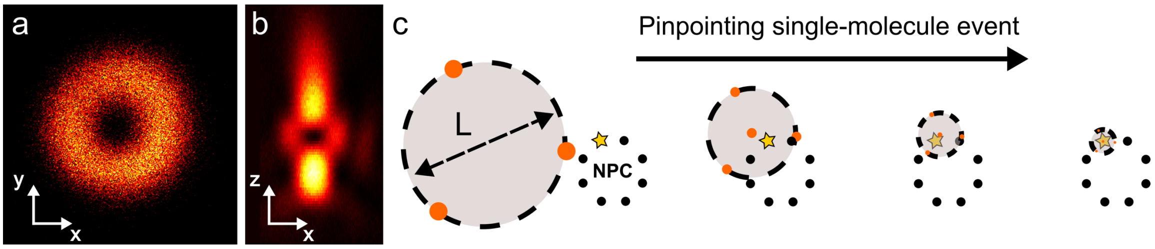


**Supplementary Figure 1. MINFLUX concepts.** On a commercial MINFLUX system the excitation laser is shaped using a spatial light modulator to make a) a donut for 2D or b) a 'bottle-beam' for 3D measurements. The laser is then scanned systematically across the region of interest (c). Once it detects a potential fluorescent signal the system homes in on the target by centering itself in an iterative process over where the best estimate for the fluorophore's position is. The search pattern diameter 'L' is gradually decreased as the event position is more precisely localized, see also Supp. Table 7.

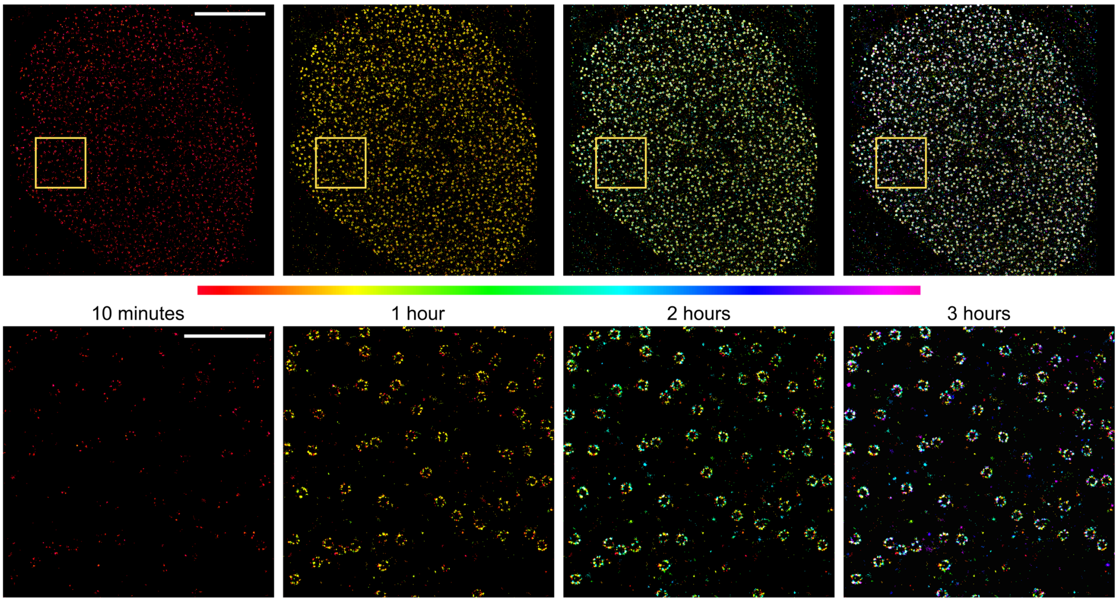


**Supplementary Figure 2. Widefield DNA-PAINT localizations at various time points. Top row:** An approximate 20x20 um region of interest, encompassing an entire cell nucleus, is imaged using a single domain antibody harboring a DNA-PAINT docking strand. With camera-based event detection the NPCs are rapidly constructed. **Bottom row:** The magnified boxed regions show that within an hour 'complete' NPC rings can already be determined. Scale bars for top row: 5 µm and for bottom row: 1 µm.

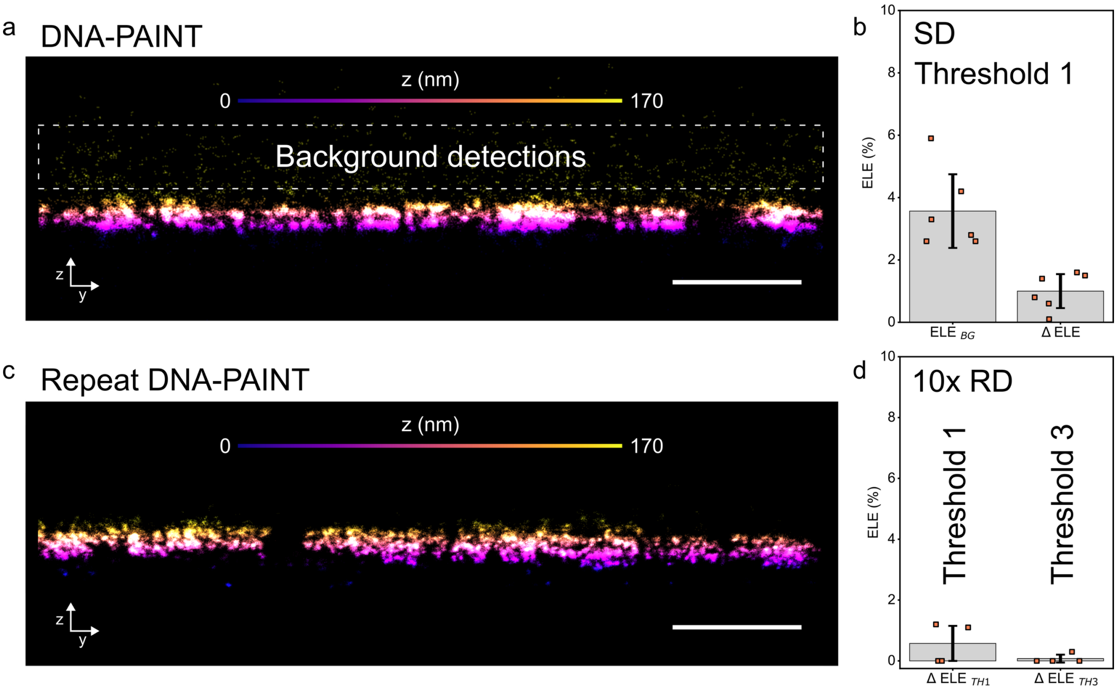


**Supplementary Figure 3. Background measurements and threshold determination. a.** A y-z view of MINFLUX 3D DNA-PAINT localizations for a medium sized region of NPCs. The background detections, highlighted in the boxed region, were used to estimate the contribution of background to our ELE measurements. **b.** Background measurements obtained ELE values of ~<4% and the effect of adding this background to NPC localizations had a marginal increase to the ELE of ~1%. **c)** Repeat DNA-PAINT experiments, conducted at approximately an order of magnitude reduced imager concentration had fewer background non-specific localizations. **d)** This resulted in a reduced contribution from background to ΔELE measurements (~0.6%) using the same threshold as for DNA-PAINT. Using a threshold relating to a minimum visit of 3 per site further reduced the contribution to ~0.1%. DNA-PAINT data N = 3, $n_{cell}$ = 6 and repeat DNA-PAINT N = 2, $n_{cell}$ = 4. Scale bar: 500 nm.

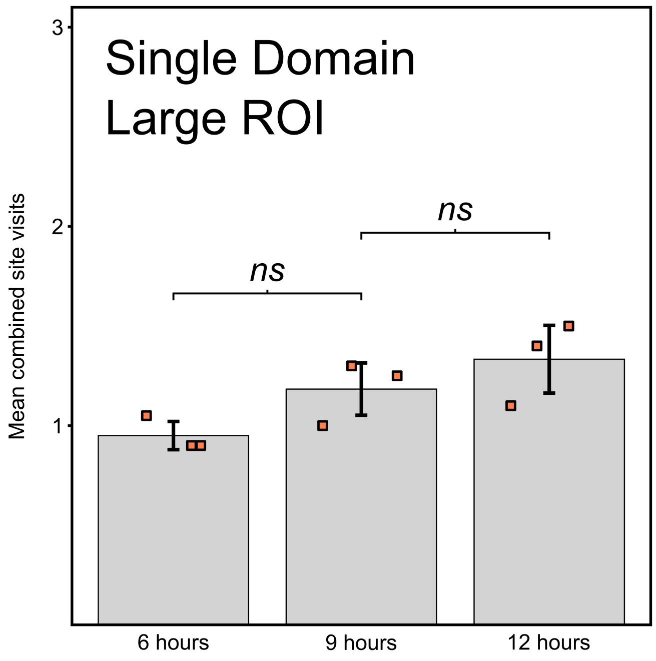


**Supplementary Figure 4. Mean combined (cytoplasmic and nucleoplasmic) site visits for single domain DNA-PAINT for a large ROI.** Imaging for longer periods of time, shown here 6, 9 and 12 hours, does not significantly improve the number of return visits obtained per site. Large ROI: N = 3, $n_{cell}$ = 3, $n_{NPC}$ = 342 (p-value>0.05).

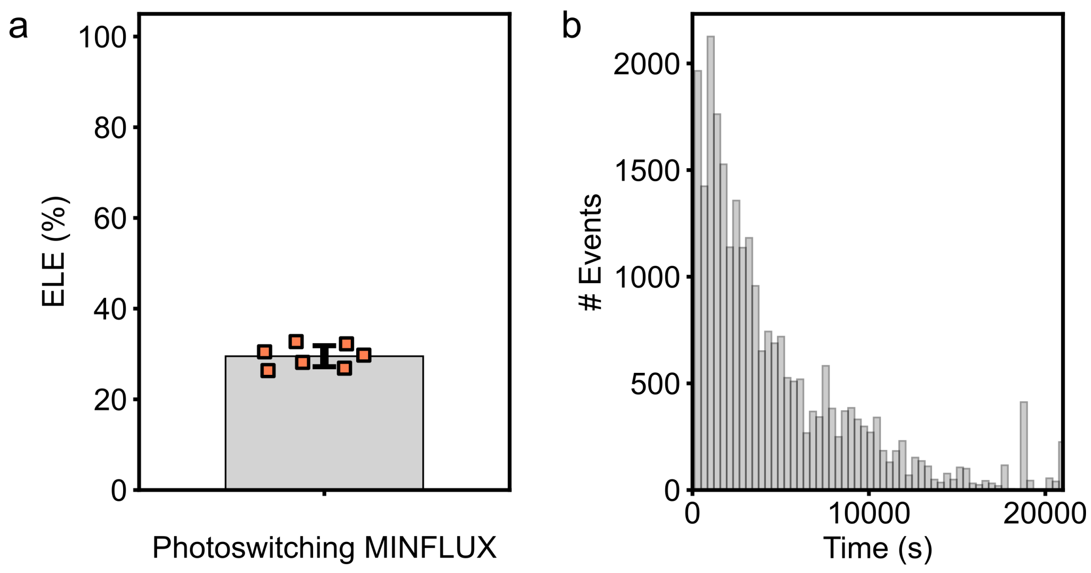


**Supplementary Figure 5. Photoswitching Alexa Fluor 647 MINFLUX data. a.** The measurements of NPCs labeled with sdAb harboring the commercial Alexa Fluor 647 dye exhibited low ELE measurements for a 3x3 'medium' sized ROI. This was credited to the rapid photobleaching resulting in fewer event detection as the experiments progressed. **b.** A histogram of events over time for a typical photoswitching MINFLUX acquisition brings a natural end to the MINFLUX experiment. Medium ROI: N = 1, $n_{cell}$ = 7, $n_{NPC}$ = 288.

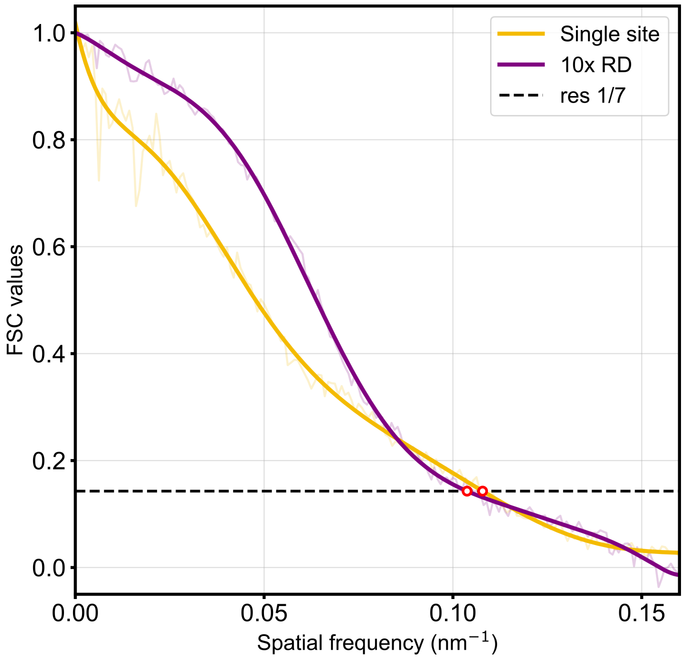


**Supplementary Figure 6. Fourier Shell Correlation (FSC) measurements for a typical 3D MINFLUX single domain DNA-PAINT (yellow) versus repeat DNA-PAINT (purple) experiment.** The 10x RD, measured under equivalent experimental conditions (duration and region size), has a similar FSC resolution measurement of 9.6 nm compared to 9.3 nm for the single domain data.

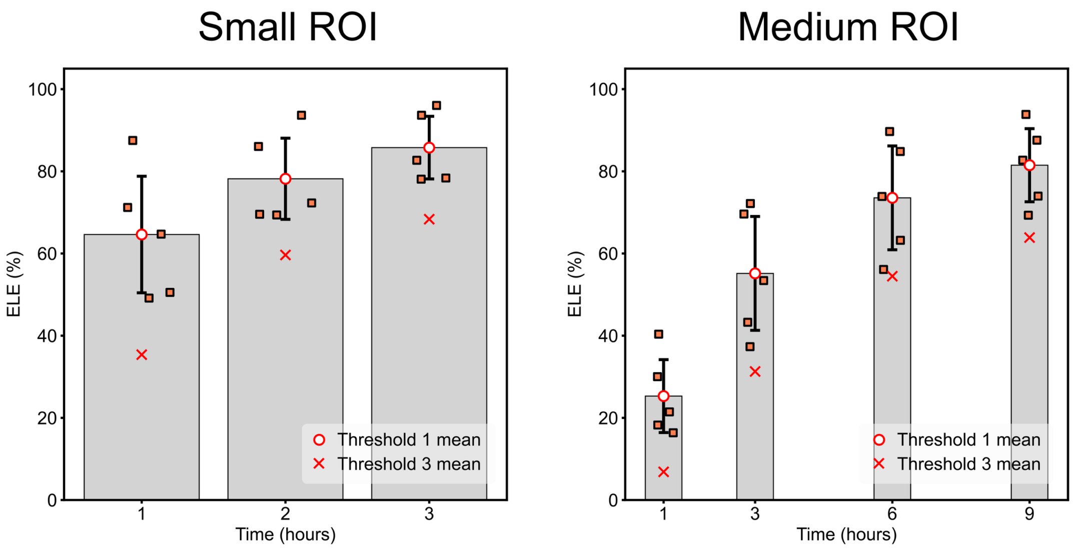


**Supplementary Figure 7. ELE measurements for repeat DNA-PAINT with a basic threshold of 1.** Using a reduced threshold stringency, the same as used for single domain DNA-PAINT experiments, the resulting ELE measurements for both small and medium ROIs are, as expected, elevated in comparison to the higher threshold of 3 visits per site where the mean value has been marked as a 'X'. These data were collected from N independent repeats, $n_{cell}$ imaged cells, and $n_{NPC}$ imaged NPCs. Small ROI: N = 3, $n_{cell}$ = 5, $n_{NPC}$ = 61, medium ROI: N =3, $n_{cell}$ = 5, $n_{NPC}$ = 177.

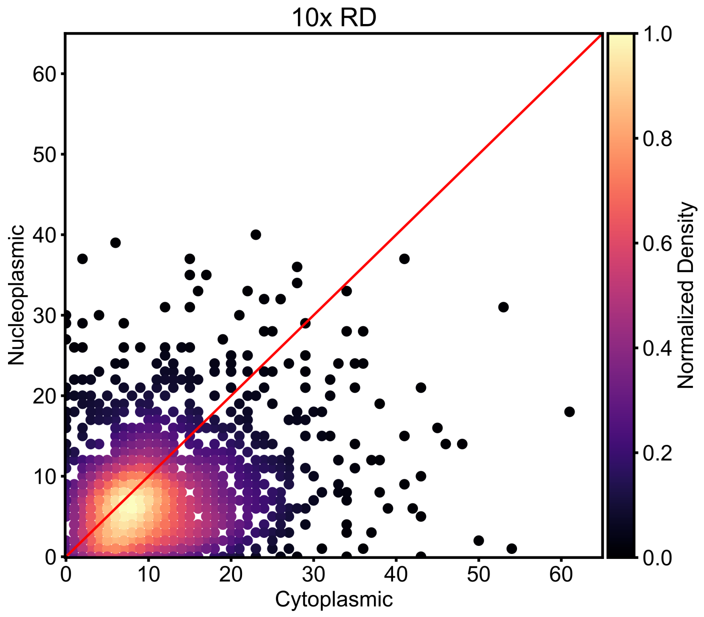


**Supplementary Figure 8. Visits to cytoplasmic sites versus nucleoplasmic sites.** In repeat DNA-PAINT experiments there was a slight increase in the number of cytosolic returns to sites compared to nucleoplasmic. N =3, $n_{cell}$ = 5, $n_{NPC}$ = 177, $n_{sites}$ = 1416.

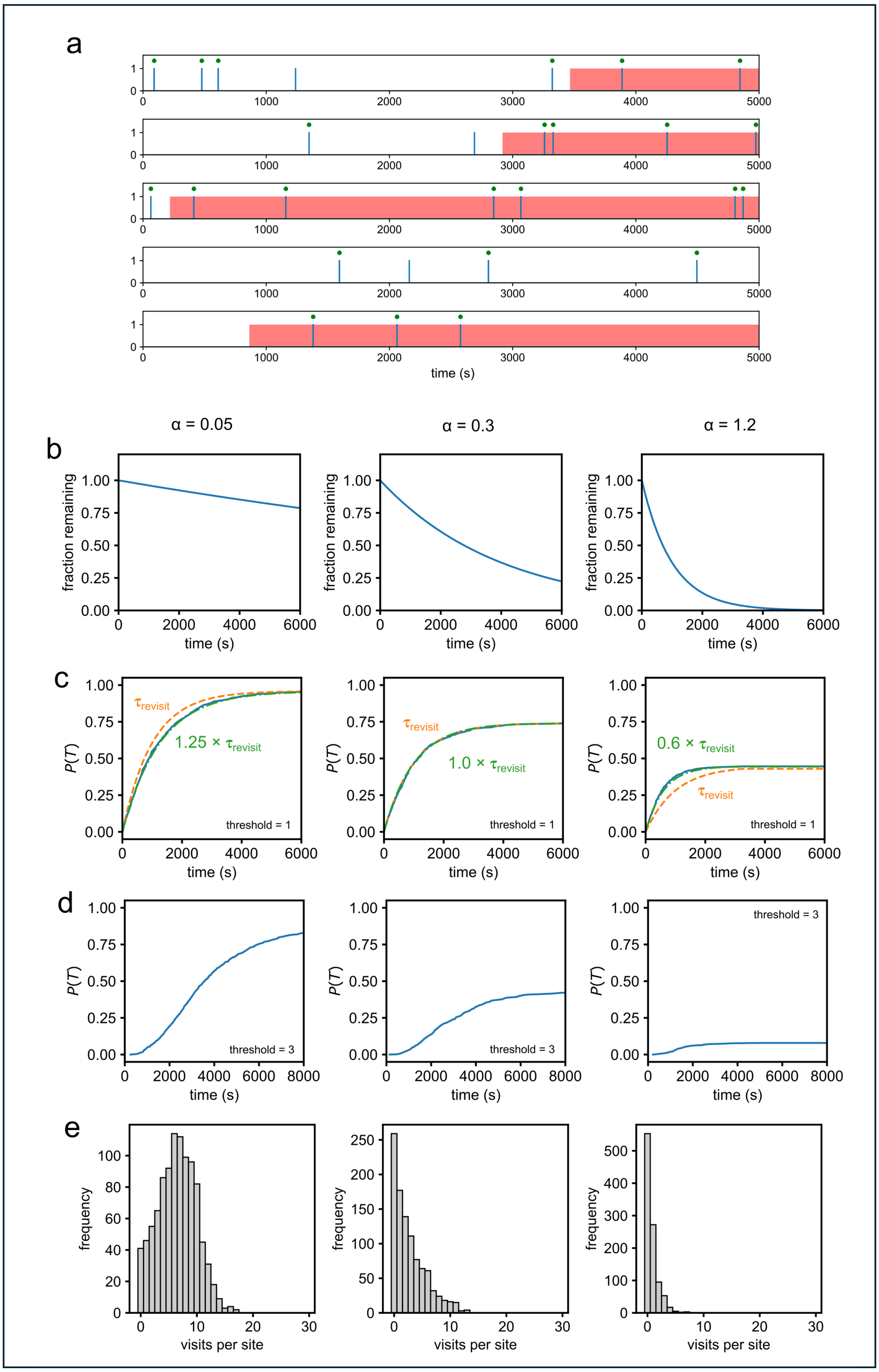

a
time (s)
b
α = 0.05
α = 0.3
α = 1.2
fraction remaining
time (s)
c
P(T)
τrevisit
1.25 × τrevisit
1.0 × τrevisit
0.6 × τrevisit
threshold = 1
d
threshold = 3
e
frequency
visits per site

**Supplementary Figure 9. A Monte Carlo model of DNA-PAINT revisits at target sites. a**. Sample trajectories from simulations with $\tau_{\text{attach}} = 0.5\ s$, $\tau_{\text{revisit}} = 1000\ s$ and a minimum detection time $t_{\text{attach,min}} = 125\ ms$. A value of 0 indicates “no attachment”, brief periods of 1 indicating “imager attachment”. Site visits longer than the minimum time required for detection are indicated with a green dot over the event. Parts of the traces overlaid in red indicate that site-loss has occurred. Attachment events in the red overlaid period are ignored (i.e. effectively no further attachment is possible once the trace is overlaid in red). **b**. Site-loss time course with $\tau_{\text{SL}} = 2.5 \cdot 10^4, 4 \cdot 10^3$ and $1 \cdot 10^3\ s$, respectively, corresponding to $\alpha = 0.05, 0.3$ and 1.2. As mentioned in the model description, we qualitatively associate the model behavior for these $\alpha$ values with “repeat DNA-PAINT-like”, “DNA-PAINT-like” and “dSTORM-like”, respectively, with the caveats as described therein. All other parameters as in **a**. **c**. Detection probability $P(\text{T})$ for a threshold of 1 visit to regard a site as “detected”. The blue curves show the fraction of detected sites as acquisition time progresses, the orange dashed lines show comparison curves with time constant $\tau_{\text{revisit}} = 1000\ s$, the green dash-dotted lines show an exponential saturation with time constant as indicated, that fit the respective time-course obtained from the simulations well. Note with little site-loss, there is a slight slowing of $P(\text{T})$ as compared to $\tau_{\text{revisit}}$, indicating that $\tau_{\text{eff}} = 1.25 \times \tau_{\text{revisit}}$ (due to some brief attachments being missed), with moderate site-loss (middle) the acceleration of saturation due to site-loss just compensates so that $\tau'_{\text{eff}} \approx \tau_{\text{revisit}}$, while with fast site-loss $\tau'_{\text{eff}} \approx 0.6 \times \tau_{\text{revisit}}$. Saturating values of detection probability are 0.95, 0.78, 0.43, respectively. **d**. Same as **c** but for a more stringent threshold of 3 visits to regard a site as “detected”. Saturating values of detection probability are then reduced to 0.85, 0.44 and 0.09, respectively. **e**. Site visit histograms for an imaging duration of $10 \times \tau_{\text{revisit}}$, i.e. 10000 s, showing the progressive change to an exponentially decaying shape as site-loss accelerates relative to characteristic site-revisit times.

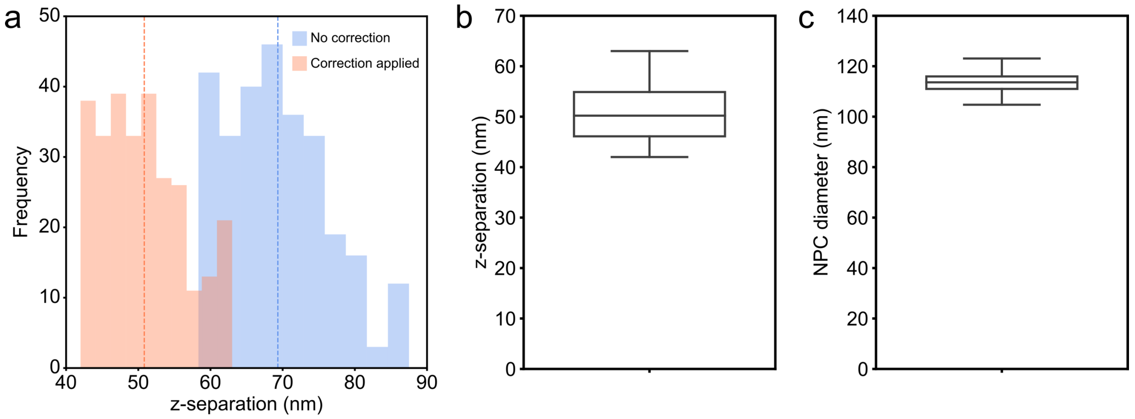


**Supplementary Figure 10. Foreshortening correction.** Imported data to the PyME visualize software had a foreshortening correction factor of 0.72 applied to all z-coordinates. **a.** Histograms for the z-separation of NPC nucleoplasmic and cytoplasmic rings with and without the foreshortening factor applied. Dashed colored lines represent the mean of the data. **b.** Boxplot representation of **b)** NPC z-separation and **c)** NPC diameter agree with expected values. Center line of boxplots shows the median value. Medium ROI: N = 4, $n_{cell}$ = 6, $n_{NPC}$ = 280.

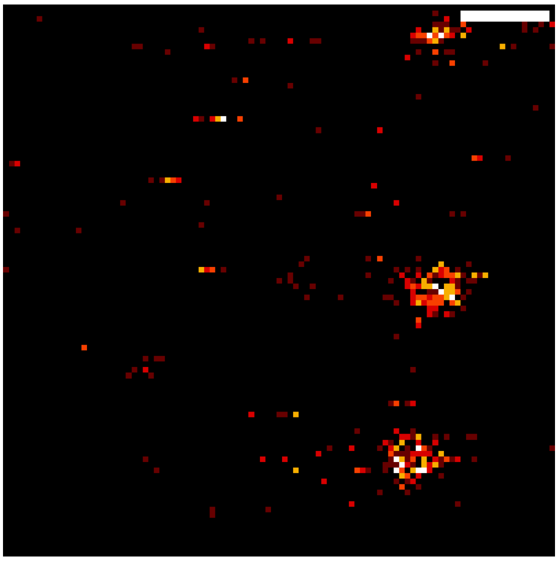

**Supplementary Media 1. Imager binding check.** Prior to commencing a MINFLUX measurement an xyt scan recorded imager binding events. The number of events observed is dependent on the labeling density of the ROI imaged. A fixed 3x3 um area measured with a 30 nm pixel size and 30 µs dwell time was used to ascertain appropriate conditions for imaging.

**Supplementary Table 1:** Single domain DNA-PAINT ELE results for different MINFLUX scan sizes measured over time.

| Scanned ROI | Hours | Mean | Std | N (repeat) | $n_{cell}$ | $n_{NPC}$ |
|---|---|---|---|---|---|---|
| **1.5 µm x 1.5 µm (Small)** | 1 | 55.9 | 13.0 | 4 | 15 | 209 |
| | 2 | 63.2 | 10.0 | | | |
| | 3 | 68.2 | 9.0 | | | |
| **3 µm x 3 µm (Medium)** | 1 | 32.7 | 9.5 | 4 | 6 | 280 |
| | 3 | 50.6 | 8.2 | | | |
| | 6 | 58.4 | 6.7 | | | |
| | 9 | 61.4 | 5.7 | | | |
| **5 µm x 5 µm (Large)** | 1 | 17.4 | 2.5 | 3 | 3 | 342 |
| | 3 | 35.8 | 2.8 | | | |
| | 6 | 47.8 | 3.6 | | | |
| | 9 | 52.3 | 4.0 | | | |
| | 12 | 54.6 | 3.5 | | | |

**Supplementary Table 2:** Single domain DNA-PAINT ELE interval measurements for different MINFLUX scan sizes.

| Scanned ROI | Time interval (hours) | Mean | Std | N (repeat) | $n_{cell}$ | $n_{NPC}$ |
|---|---|---|---|---|---|---|
| **1.5 µm x 1.5 µm (Small)** | 0-1 | 55.9 | 13.0 | 4 | 15 | 209 |
| | 1-2 | 32.3 | 8.0 | | | |
| | 2-3 | 21.4 | 4.8 | | | |
| **3 µm x 3 µm (Medium)** | 0-1 | 32.7 | 9.5 | 4 | 6 | 280 |
| | 1-2 | 24.3 | 4.6 | | | |
| | 2-3 | 15.8 | 2.3 | | | |
| | 5-6 | 9.8 | 1.5 | | | |
| | 8-9 | 6.6 | 1.4 | | | |
| **5 µm x 5 µm (Large)** | 0-1 | 17.4 | 2.5 | 3 | 3 | 342 |
| | 1-2 | 14.4 | 1.1 | | | |
| | 2-3 | 14.0 | 2.0 | | | |
| | 5-6 | 8.9 | 1.6 | | | |
| | 8-9 | 6.7 | 1.2 | | | |
| | 11-12 | 6.2 | 0.9 | | | |

**Supplementary Table 3:** Alexa Fluor 647 ELE results for different MINFLUX scan sizes measured over time.

| Scanned ROI | Hours | Mean | Std | N (repeat) | $n_{cell}$ | $n_{NPC}$ |
|---|---|---|---|---|---|---|
| **3 µm x 3 µm (Medium)** | 1 | 23.4 | 4.5 | 1 | 7 | 288 |
| | Max | 29.5 | 2.3 | | | |

**Supplementary Table 4:** Repeat DNA-PAINT ELE results for different MINFLUX scan sizes measured over time using an NPC analysis threshold value of 3.

| Scanned ROI | Hours | Mean | Std | N (repeat) | $n_{cell}$ | $n_{NPC}$ |
|---|---|---|---|---|---|---|
| **1.5 µm x 1.5 µm (Small)** | 1 | 35.4 | 14.9 | 3 | 5 | 61 |
| | 2 | 59.6 | 12.8 | | | |
| | 3 | 68.4 | 7.7 | | | |
| **3 µm x 3 µm (Medium)** | 1 | 6.9 | 5.6 | 3 | 5 | 177 |
| | 3 | 31.3 | 13.7 | | | |
| | 6 | 54.5 | 12.3 | | | |
| | 9 | 63.9 | 11.2 | | | |

**Supplementary Table 5:** Repeat DNA-PAINT ELE results for different MINFLUX scan sizes measured over time using an NPC analysis threshold value of 1.

| Scanned ROI | Hours | Mean | Std | N (repeat) | $n_{cell}$ | $n_{NPC}$ |
|---|---|---|---|---|---|---|
| **1.5 µm x 1.5 µm (Small)** | 1 | 64.6 | 14.2 | 3 | 5 | 61 |
| | 2 | 78.2 | 9.9 | | | |
| | 3 | 85.8 | 7.6 | | | |
| **3 µm x 3 µm (Medium)** | 1 | 25.3 | 8.9 | 3 | 5 | 177 |
| | 3 | 55.2 | 13.9 | | | |
| | 6 | 73.6 | 12.6 | | | |
| | 9 | 81.5 | 8.9 | | | |

**Supplementary Table 6:** Repeat DNA-PAINT ELE interval measurements for different MINFLUX scan sizes using an NPC analysis threshold value of 3.

| Scanned ROI | Hours | Mean | Std | N (repeat) | $n_{cell}$ | $n_{NPC}$ |
|---|---|---|---|---|---|---|
| **1.5 µm x 1.5 µm (Small)** | 0-1 | 35.4 | 14.9 | 3 | 5 | 61 |
| | 1-2 | 36.7 | 10.8 | | | |
| | 2-3 | 33.1 | 10.3 | | | |
| **3 µm x 3 µm (Medium)** | 0-1 | 6.9 | 5.6 | 3 | 5 | 177 |
| | 1-2 | 8.4 | 5.1 | | | |
| | 2-3 | 8.3 | 4.4 | | | |
| | 5-6 | 7.8 | 3.4 | | | |
| | 8-9 | 6.7 | 2.4 | | | |

**Supplementary Table 7:** 3D MINFLUX imaging sequence parameters.

| Itr | Pattern | L (nm) | patGeoFactor | phtLimit | patDwellTime | patRepeat | pwrFactor | ccrLimit (cfr check) | bgcThreshold |
|---|---|---|---|---|---|---|---|---|---|
| 0 | hexagon | 288 | 0.8 | 160 | 0.001 | 1 | 1 | -1 | 15000 |
| 1 | zline | 288/1440 | 0.8/4 | 400 | 0.001 | 1 | 1 | -1 | 15000 |
| 2 | square | 288 | 0.8 | 100 | 0.001 | 5 | 1 | -1 | 10000 |
| 3 | zline2 | 288 | 0.8 | 50 | 0.001 | 5 | 1 | -1 | 10000 |
| 4 | square | 151.2 | 0.42 | 67 | 0.001 | 5 | 2 | 0.9 | 10000 |
| 5 | zline2 | 151.2 | 0.42 | 33 | 0.001 | 5 | 2 | -1 | 10000 |
| 6 | square | 75.6 | 0.21 | 67 | 0.001 | 5 | 4 | 0.8 | 10000 |
| 7 | zline2 | 75.6 | 0.21 | 33 | 0.001 | 5 | 4 | -1 | 10000 |
| 8 | square | 39.6 | 0.11 | 100 | 0.001 | 5 | 6 | -1 | 10000 |
| 9 | zline2 | 39.6 | 0.11 | 50 | 0.001 | 5 | 6 | -1 | 10000 |
